# OBERON3 and SUPPRESSOR OF MAX2 1-LIKE proteins form a regulatory module specifying phloem identity

**DOI:** 10.1101/2019.12.21.885863

**Authors:** Eva-Sophie Wallner, Nina Tonn, Dongbo Shi, Laura Luzzietti, Friederike Wanke, Pascal Hunziker, Yingqiang Xu, Ilona Jung, Vadir Lopéz-Salmerón, Michael Gebert, Christian Wenzl, Jan U. Lohmann, Klaus Harter, Thomas Greb

## Abstract

Spatial specificity of cell fate decisions is central for organismal development. The phloem tissue mediates long-distance transport of energy metabolites along plant bodies and is characterized by an exceptional degree of cellular specialization. How a phloem-specific developmental program is implemented is, however, unknown. Here we reveal that the ubiquitously expressed PHD-finger protein OBE3 forms a central module with the phloem-specific SMXL5 protein for establishing phloem identity in *Arabidopsis thaliana*. By protein interaction studies and phloem-specific ATAC-seq analyses, we show that OBE3 and SMXL5 proteins form a complex in nuclei of phloem stem cells where they establish a phloem-specific chromatin profile. This profile allows expression of *OPS*, *BRX*, *BAM3*, and *CVP2* genes acting as mediators of phloem differentiation. Our findings demonstrate that OBE3/SMXL5 protein complexes establish nuclear features essential for determining phloem cell fate and highlight how a combination of ubiquitous and local regulators generate specificity of developmental decisions in plants.

## Introduction

Growth and body shape of multicellular organisms largely depend on a functional long-distance transport of energy metabolites to fuel stem cell activity. In vascular plants, sugars are photosynthetically produced in source organs, such as leaves, and delivered via the phloem to sink organs where they are allocated to storage tissues or stem cell niches, such as the root apical meristem (RAM) (Oparka and Turgeon 1999; De Schepper et al. 2013). The dividing stem cells of the RAM are located next to a mostly dormant organizer, known as quiescent center (QC) (van den Berg et al. 1997). These stem cells divide and differentiate in a strictly controlled manner to give rise to two phloem poles which ensure a steady energy supply to the RAM during root growth (Rodriguez-Villalon et al. 2014; Wallner et al. 2017). One phloem pole comprises a protophloem and a metaphloem strand, each forming a sieve element (SE) and a companion cell (CC) lineage (Lucas et al. 2013). During differentiation, SEs degrade most of their organelles to build connected sieve tubes for intracellular allocation of sugars, hormones, proteins and RNAs (Furuta et al. 2014). This is why functional SEs are metabolically sustained by CCs via intercellular channels named plasmodesmata (Ross-Elliott et al. 2017). Underlining the importance of the phloem, defects in protophloem development impair root growth, possibly, as a consequence of RAM starvation (Depuydt et al. 2013; Rodriguez-Villalon et al. 2014).

Due to the remarkable transition of phloem stem cells to cells holding an extreme degree of specialization, gaining insights into phloem formation and identifying its molecular regulators is highly instructive for our general understanding of cell fate regulation and differentiation (Marhava et al. 2018; Gujas et al. 2020; Marhava et al. 2020). Moreover, due to the importance of the phloem for plant growth and physiology, revealing mechanisms of phloem formation holds great promises for crop production and may increase our understanding of plant evolution and of the adaptation to environmental conditions (López-Salmerón et al. 2019). Importantly, although several genes, including *ALTERED PHLOEM DEVLEOPMENT* (*APL*), *OCTOPUS* (*OPS*), *BREVIS RADIX* (*BRX*), *BARELY ANY MERISTEM3* (*BAM3*), and *COTYLEDON VASCULAR PATTERN2* (*CVP2*) have been identified to regulate different aspects of phloem formation (Bonke et al. 2003; Depuydt et al. 2013; Rodriguez-Villalon et al. 2014; Anne et al. 2015; Rodriguez-Villalon et al. 2015; Hazak et al. 2017; Marhava et al. 2018), those genes seem to act downstream of phloem specification leaving the question open of how a phloem-specific developmental program is initiated.

Recently, we revealed a central role of the SUPPRESSOR OF MAX2 1-LIKE (SMXL) protein family members SMXL3, SMXL4 and SMXL5 in phloem formation (Wallner et al. 2017; Wallner et al. 2020). SMXL proteins are well-conserved nuclear localized developmental regulators and, in *Arabidopsis thaliana* (Arabidopsis), form a protein family of eight members sub-divided into different sub-clades based on phylogeny and function (Zhou et al. 2013; Soundappan et al. 2015; Liang et al. 2016; Wallner et al. 2016; Walker and Bennett 2017). Among those, SMXL6, SMXL7, and SMXL8 are proteolytic targets of the strigolactone signaling pathway which bind directly to promoter regions of downstream target genes and, thereby, repress their transcription (Soundappan et al. 2015; Wang et al. 2015; Liang et al. 2016; Ma et al. 2017; Wang et al. 2020). In comparison, SMXL3/4/5 proteins act independently from strigolactone signaling as central regulators of phloem formation (Wallner et al. 2017; Wu et al. 2017; Cho et al. 2018). Their redundant and dose-dependent functions become obvious in double and triple mutants which are completely deprived of protophloem formation within the RAM resulting in root growth termination a few days after germination (Wallner et al. 2017). Despite their fundamental role in phloem formation the mechanism of SMXL3/4/5 protein action remained obscure.

In contrast to SMXL proteins whose activity is spatially highly restricted, OBERONs (OBEs) are a family of four ubiquitously expressed, nuclear-localized proteins essential for tissue specification and meristem maintenance starting from the earliest stages of embryo development (Saiga et al. 2008; Thomas et al. 2009; Saiga et al. 2012; Lin et al. 2016). This role is reflected by mutants deficient for either of the two *OBE* sub-families which are embryo lethal (Saiga et al. 2008; Thomas et al. 2009; Saiga et al. 2012). In the shoot apical meristem (SAM), *OBE3* (also known as *TITANIA1* (*TTA1*)), interacts genetically with the homeobox transcription factor gene *WUSCHEL* (*WUS*) in stem cell regulation (Lin et al. 2016). Additionally, *OBE1* and *OBE2* are associated with vascular patterning in the embryo (Thomas et al. 2009). Interestingly, OBEs carry a highly conserved plant homeodomain (PHD)-finger domain known to bind di- and trimethylated histone H3 which allows recruitment of chromatin remodeling complexes and transcription factors (Sanchez and Zhou 2011). Indeed, OBE proteins show chromatin binding and remodeling activities important for root initiation during embryogenesis (Saiga et al. 2008; Saiga et al. 2012). Taken together, OBEs have versatile roles associated with cell fate regulation in plants (Saiga et al. 2008; Thomas et al. 2009; Saiga et al. 2012; Lin et al. 2016) but, as for SMXL proteins, their specific roles in distinct tissues and their mode of action is unknown.

Here, we report that OBE3 and SMXL5 proteins physically interact forming a functional unit during protophloem formation in the RAM. We provide evidence that SMXL5 and OBE3 proteins are instrumental for the establishment of a phloem-specific chromatin configuration and for the expression of phloem-associated regulators. By characterizing the *SMXL3/4/5- OBE3* interaction and of phloem-specific chromatin conformation we provide insights into molecular mechanisms of cell specification and the establishment of a highly specialized and central plant tissue.

## Results

### *SMXL4* and *SMXL5* promote expression of early phloem markers

To map the function of the *SMXL4* and *SMXL5* genes within the process of phloem formation, we introgressed a series of developmental markers visualizing early steps of phloem formation (Rodriguez-Villalon 2016) into the *smxl4;smxl5* double mutants showing severe defects in protophloem formation (Wallner et al. 2017). Analysis of root tips two days after germination when the overall anatomy of the *smxl4;smxl5* RAM is comparable to wild type RAMs (Wallner et al. 2017), showed that *OPS:OPS-GFP, BRX:BRX-CITRINE*, *BAM3:BAM3-CITRINE*, or *CVP2:NLS-VENUS* marker activities (Rodriguez-Villalon et al. 2014) were reduced or not detectable in *smxl4;smxl5* plants (Figure 1, A-H). This reduction was found along the entire strand of the developing protophloem and included SE-procambium stem cells located immediately proximal to the quiescent center (QC). In these founder cells of the phloem lineage, we observed accumulation of OPS-GFP and BRX-CITRINE fusion proteins in wild type which was hardly detectable in *smxl4;smxl5* double mutants (Figure 1, I-P). In contrast to markers associated with early stages of phloem development, activity of the *APL* promoter marking differentiating SEs and CCs in wild type (Bonke et al. 2003) was not detectable in root tips of *smxl4;smxl5* mutants (Supplementary Figure 1). These observations argued for an early stem cell-associated role of *SMXL4* and *SMXL5* in establishing a general phloem-specific developmental program including *OPS*, *BRX*, *BAM3*, and *CVP2* gene activities. Supporting this conclusion, *smxl4;smxl5;bam3* triple mutants showed root growth defects similar to *smxl4;smxl5* double mutants and stimulating the BAM3 pathway by CLE45 treatments had no effect on *smxl4;smxl5* roots (Supplementary Figure 1). Together with the reduced *BAM3* reporter activity (Figure 1, E,F), this indicated that the phloem-specific BAM3 pathway which counteracts phloem development (Depuydt et al. 2013) is less active in *smxl4;smxl5* mutants and not causing the developmental defects observed in *smxl4;smxl5*.

**Figure 1:**
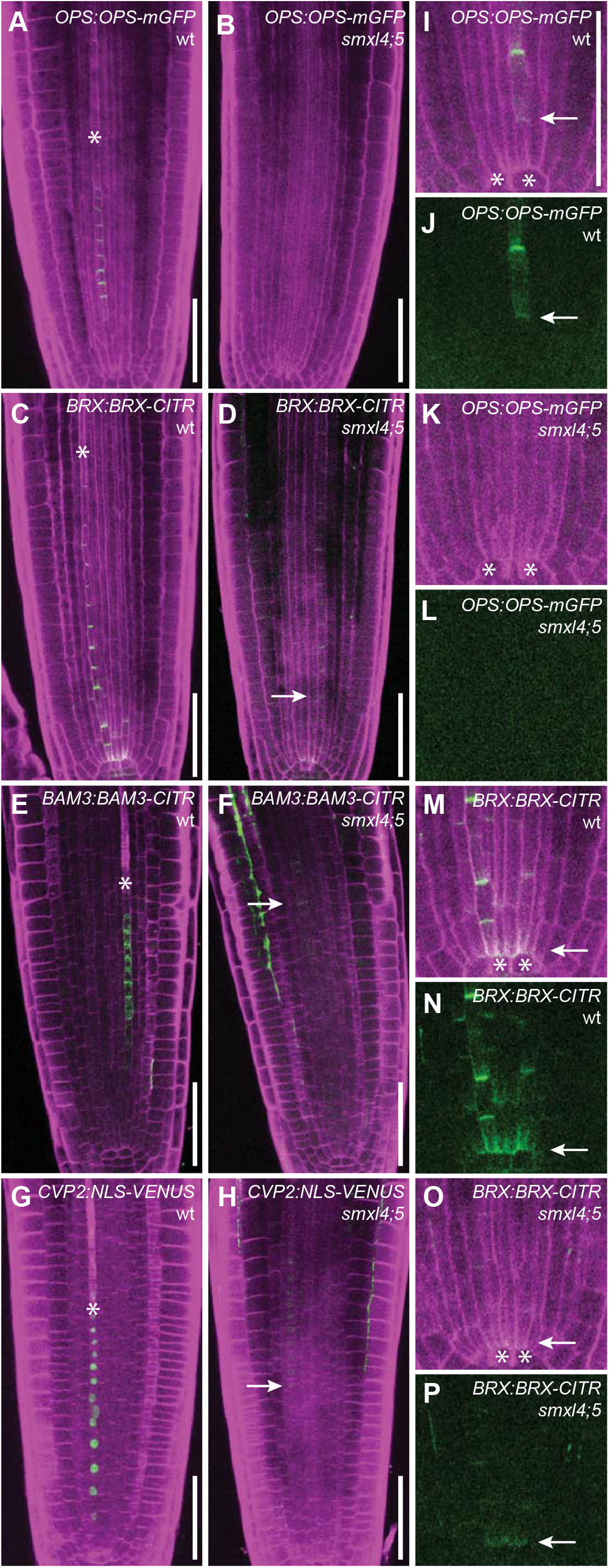
Phloem-related genes are less active in *smxl4;smxl5* mutants. **(A – H)** Comparison of *OPS:OPS-GFP* (A, B), *BRX:BRX-CITRINE* (C, D), *BAM3:BAM3-CITRINE*, and *CVP2:NLS-VENUS* (G, H) reporter activities in wild type (A, C, E, G) and *smxl4;smxl5* double mutants (B, D, F, H). Asterisks depict the first differentiating SE. Arrows point to earliest detectable reporter activities. Scale bars represent 50 µm. Note that the fluorescent signal in the root cortex visible in F is autofluorescence due to tissue disruption. **(I – P)** *OPS:OPS-GFP* (I-L) and *BRX:BRX-CITRINE* (M-P) close to the QC in wild type (I, J, M, N) and *smxl4;smxl5* mutants (K, L, O, P). In J, L, N, and P, fluorescent signals are depicted without counterstaining. Asterisks indicate QC cells. Arrows point to earliest detectable reporter activities. Scale bar in (I) represents 50 µm. Same magnification in I-P.

### Activity of *SMXL* genes upstream of *OPS* and *BRX* is required for phloem formation

To challenge the idea of an early role of *SMXL5*, we tested the capacity of *SMXL5* to restore protophloem formation when expressed under the control of promoters active during different phases of protophloem development (Rodriguez-Villalon 2016). Root length served as a fast and efficient read-out for phloem defects (Depuydt et al. 2013; Wallner et al. 2017). Supporting the need of *SMXL5* activity during early phases of phloem development, reduced root length and impaired SE formation usually found in *smxl4;smxl5* mutants was not observed when they expressed *SMXL5* under the control of the early *OPS*, *BAM3*, or *CVP2* promoters (Figure 2A, Supplementary Figure 1). In contrast, driving *SMXL5* expression by the late *APL* promoter did not restore of root length or SE formation (Figure 2A, Supplementary Figure 1).

**Figure 2:**
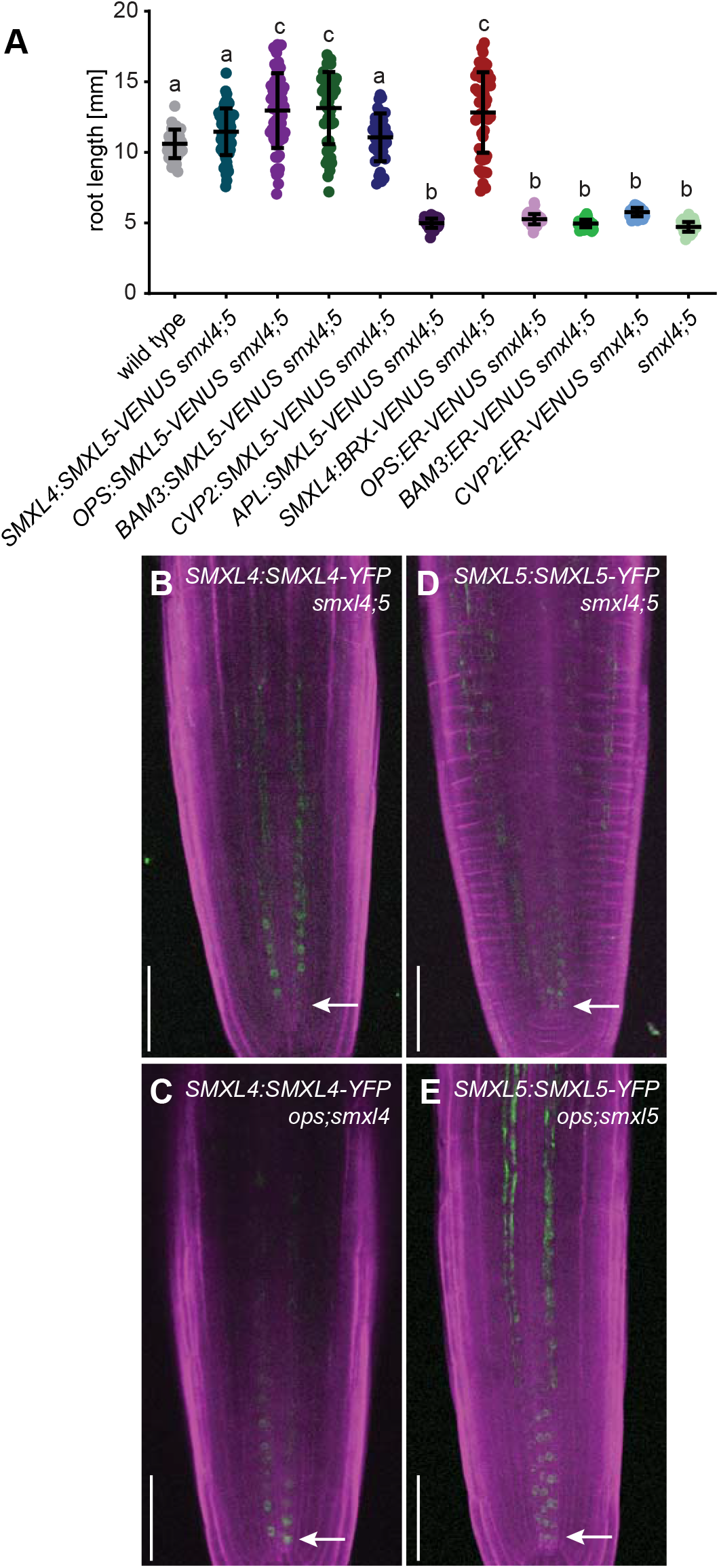
Analysis of interaction between *SMXL4* and *SMXL5* with other phloem regulators. (**A**) Root length of five day-old plants. n = 37–75. Statistical groups determined by one-way ANOVA and post-hoc Tukey’s test (95 % CI). Shown is one representative experiment of three repetitions. **(B – E)** Comparison of *SMXL4:SMXL4-YFP* and *SMXL5:SMXL5-YFP* reporter activities in *smxl4;smxl5* and *smxl;ops* mutants. Arrows indicate the signals closest to the QC. Scale bars represent 50 µm.

To see whether the reduced activity of regulators like *BRX* is causative for reduced root length of *smxl4;smxl5* double mutants, we expressed *BRX-VENUS* in *smxl4;smxl5* mutant backgrounds under the control of the protophloem-specific *SMXL4* promoter (Supplementary Figure 1F) (Wallner et al. 2017). Indeed, root length of *SMXL4:BRX-VENUS/smxl4;smxl5* lines was comparable to the length of wild type roots (Figure 2A) indicating that *BRX* acts downstream of *SMXL4* and that reduced *BRX* activity is one reason for disturbed phloem development in *smxl4;smxl5* mutants. Further supporting a role of the *SMXL4* and *SMXL5* genes upstream of *OPS*, visualization of SMXL4 and SMXL5 proteins in *OPS*-deficient backgrounds by respective reporters (Wallner et al. 2017) did neither reveal a reduced level nor altered localization of SMXL4 or SMXL5 proteins in protophloem cells (Figure 2, B-E). This suggested that in contrast to a positive effect of *SMXL4* and *SMXL5* on the activity of *OPS* and *BRX* genes (Figure 1), *OPS* was not important for stimulating *SMXL4* or *SMXL5* activity and that *OPS* and *SMXL* genes function at distinct steps during phloem formation.

### *SMXL* genes act on different steps of phloem formation than *OPS* and *BRX*

Our interpretation that *SMXL* genes on the one side and *OPS* and *BRX* on the other side act on different steps of phloem development, was confirmed by investigating their genetic interaction. The OPS protein is required for SE formation in the protophloem by counteracting the BAM3/CLAVATA3/ESR-related 45 (CLE45) pathway (Breda et al. 2019). Due to enhanced activity of the BAM3/CLE45 pathway, *ops* mutants develop ‘gap cells’ within protophloem strands in which SE establishment fails (Truernit et al. 2012; Rodriguez-Villalon et al. 2014). Interestingly, root length of *ops;smxl5* double mutants was similarly reduced as in *smxl4;smxl5* double mutants although *smxl5* single mutants were not affected and *ops* single mutants were very variable in this regard (Figure 3A). This finding suggested an additive effect of *SMXL* and *OPS*-dependent pathways on phloem formation. Indeed, when phloem development was carefully analyzed in the respective mutant backgrounds, we observed a variation of phloem defects in *ops* single mutants ranging from the appearance of gap cells to the complete absence of SEs in a small fraction of plants (Figure 3B, Supplementary Figure 2). In comparison, 60 % of the *ops;smxl5* mutants displayed complete SE deficiency demonstrating that both genes contribute to a robust phloem development. A similar trend was observed for *brx;smxl5* double mutants. Like *OPS*, the *BRX* gene ensures continuous SE formation, in this case, by downregulation of *BAM3* transcription and steepening the auxin gradient in developing phloem cells (Scacchi et al. 2009; Depuydt et al. 2013; Marhava et al. 2018; Marhava et al. 2020). Similar to *ops;smxl5* double mutants, *brx;smxl5* plants developed shorter roots than *brx* and *smxl5* single mutants and largely failed to differentiate SEs (Figure 3A, Supplementary Figure 3). Importantly, in those *ops;smxl5* plants developing SEs, gap cell formation was comparable to *ops* and *brx* single mutants (Figure 3B, Supplementary Figures 1). These observations suggested that, although all three genes are important for stable SE formation, *SMXL5* and *OPS*/*BRX* genes play roles at different steps during phloem formation with *SMXL5* acting upstream.

**Figure 3:**
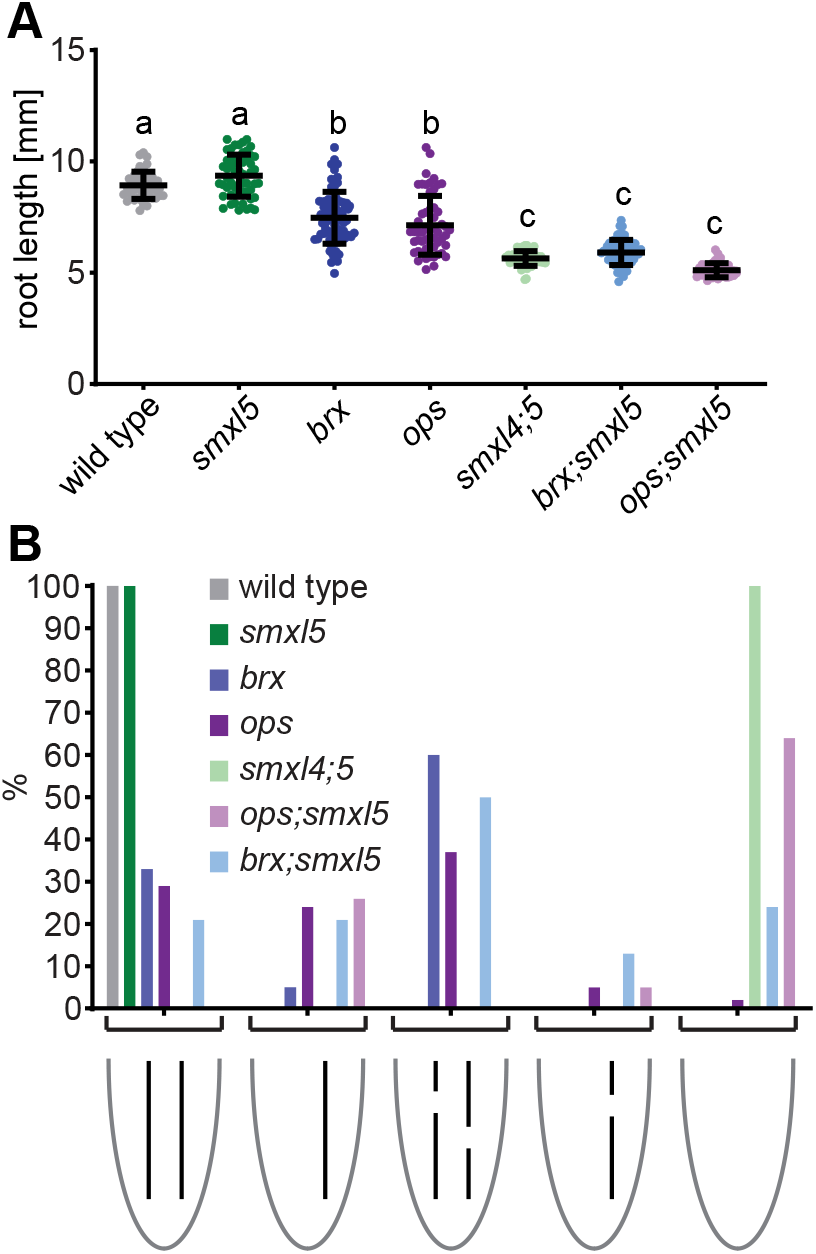
Genetic interaction of *SMXL5* with *OPS* and *BRX*. **(A)** Root length of five day-old wild type (WT), *smxl5*, *brx*, *ops*, *smxl4;smxl5*, *brx;smxl5*, and *ops;smxl5* plants. n = 38–76 for each group. Statistical groups determined by one-way ANOVA and post-hoc Tukey’s HSD test (95 % CI). Shown is one representative experiment of three repetitions. **(B)** Phenotypic characterization of phloem development of two day-old wild type, *smxl5*, *brx*, *ops*, *smxl4;smxl5*, *brx;smxl5*, and *ops;smxl5* plants. n = 97–129 for each group.

### SMXL5 proteins interact with OBE3 proteins in nuclei of plant cells

To find indications for how SMXL proteins fulfil their early role, we isolated interacting proteins by a Yeast-Two-Hybrid-based screen of a cDNA expression library generated from *Arabidopsis* seedlings (Legrain et al. 2001; Cifuentes-Esquivel et al. 2013) using the full length SMXL5 protein as a bait. After testing 84 million individual protein-protein interactions, we identified OBE3 as one out of five ‘very high confident’ candidate interactors with 24 isolated independent cDNA clones (Supplementary Figure 4). After confirming the yeast-based interaction of OBE3 with SMXL5 in independent experiments (Figure 4A), we tested whether both proteins also interact *in planta*. To this end, we transiently expressed SMXL5 fused to a triple human influenza hemagglutinin (HA) affinity tag and OBE3 fused to a six fold c-Myc epitope tag in *Nicotiana benthamiana* (*N. benthamiana*) leaves under the control of the Cauliflower Mosaic Virus (CaMV) *35S* promoter (Benfey and Chua 1990). In raw protein extracts before (‘input’) and after (‘unbound’) immunoprecipitation (IP) using HA-affinity beads and in the precipitate itself (‘IP: α HA’), the SMXL5-3xHA protein was detected with the expected size of approximately 120 kDa in Western analyses (Figure 4B). Importantly, the 6xMyc-OBE3 fusion protein co-immunoprecipitated with the SMXL5-3xHA protein and did not show unspecific binding to the HA-affinity beads, indicating that SMXL5-3xHA and 6xMyc- OBE3 proteins interacted in plant cells.

**Figure 4:**
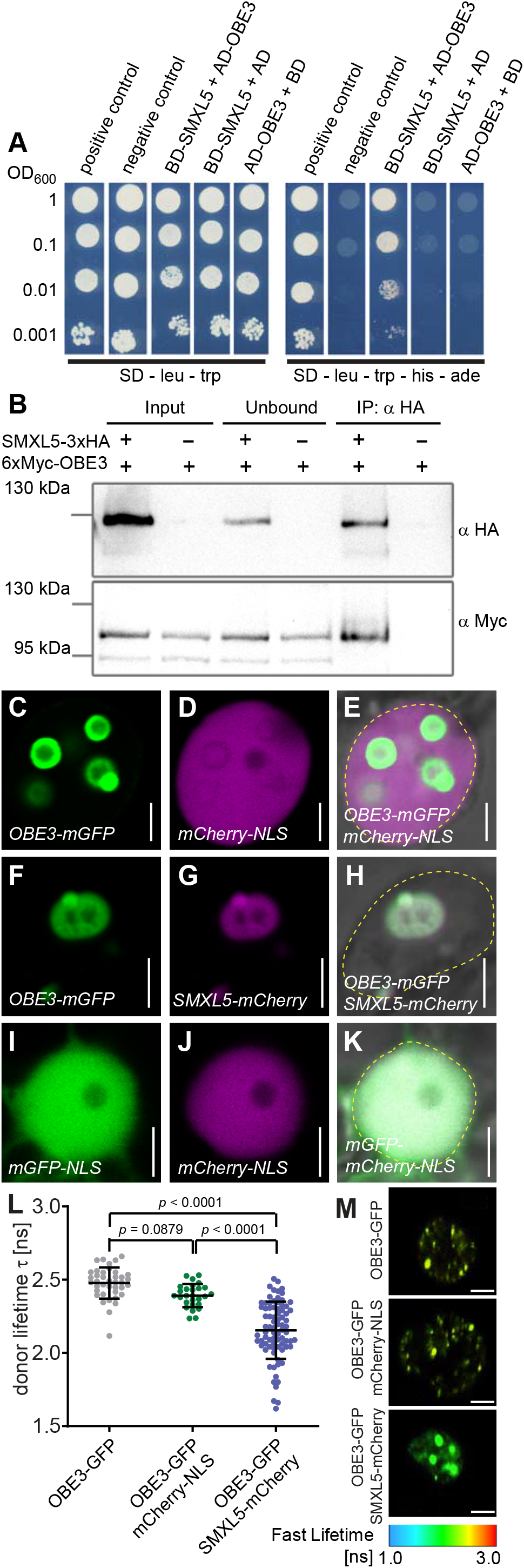
SMXL5 and OBE3 proteins interact. **(A)** The SMXL5 protein was expressed in yeast fused to the GAL4 DNA binding domain (BD) and OBE3 fused to the GAL4 activation domain (AD). All strains contained AD- and BD- expressing plasmids either alone or fused with SMXL5, OBE3 or control proteins, respectively. Growth on SD-leu-trp indicate the presence of both plasmids, growth on SD-leu-trp-his-ade medium indicate the presence of plasmids and protein interaction. **(B)** Interaction of SMXL5-3xHA and 6xMyc-OBE3 proteins by co-immunoprecipitation (co-IP) and subsequent Western blot analysis after transient overexpression in *N. benthamiana*. ‘Input’ represents unprocessed protein extracts, ‘unbound’ show proteins that remained in the extract after IP and ‘IP: α HA’ depicts samples after immunoprecipitation by α-HA-beads. Western blots were probed by α HA or α Myc antibodies, respectively. Signals revealed an expected SMXL5-3xHA protein size of approximately 120 kDa. The size of the detected 6xMyc-OBE3 protein (∼100 kDa) exceeded the expected size of 92 kDa. **(C – K)** Fluorescent signals and bright field images of epidermal *N. benthamiana* nuclei transiently co-expressing OBE3-mGFP/mCherry (C-E), OBE3-mGFP/SMXL5-mCherry (F-H) and mGFP-NLS/mCherry-NLS (I-K). The dashed yellow line indicates the outlines of nuclei in merged images (E, H, K). Scale bars represent 5 µm. **(L)** FRET-FLIM analysis of transiently transformed *N. benthamiana* epidermal leaf cells expressing the OBE3-mGFP donor without an mCherry acceptor or in the presence of NLS- mCherry or SMXL5-mCherry. Error bars indicate standard deviation. Data were derived from three biological replicates with n = 27–80. To test for homogeneity of variance, a Brown- Forsythe test was applied. As the variances are not homogenous a Wilcoxon / Kruskal-Wallis test was carried out followed by a Dunn post hoc analysis. **(M)** Heat maps of representative nuclei used for FLIM measurements. The donor lifetimes of OBE3-mGFP are color-coded according to the scale at the bottom. Size bars represent 4 µm.

To compare the sub-cellular localizations of SMXL5 and OBE3 proteins, we transiently expressed the SMXL5 protein fused to monomeric Cherry (SMXL5-mCherry) together with the OBE3 protein fused to monomeric GFP (OBE3-mGFP) again in *N*. *benthamiana* leaves. Initially, nuclear localization of the OBE3 protein was confirmed by co-expressing OBE3-GFP with mCherry fused to a nuclear localization signal (mCherry-NLS). Interestingly, while the mCherry-NLS signal was homogenously distributed within the nucleus, the OBE3-mGFP protein appeared in nuclear subdomains (Figure 4, C-E). Co-expression of SMXL5-mCherry and OBE3-GFP revealed a co-localization of both proteins within these domains (Figure 4, F-H) which were distinct from the whole nucleus highlighted by an *mGFP-mCherry-NLS* fusion protein expressed under the control of the *ubiquitin 10* (*UBI10*) promoter (Figure 4, I-K). Next, we evaluated our yeast two-hybrid and co-immunoprecipitation data by performing Förster resonance energy transfer (FRET)-fluorescence lifetime imaging microscopy (FLIM) analysis as an *in planta* assay for protein-protein association. In transiently transformed *N. benthamiana* epidermal leaf cells, FRET-FLIM analysis detected a significant decrease in the lifetime of the donor OBE3-mGFP fusions in the nucleus when co-expressed with SMXL5-mCherry (Figure 4L, M). In contrast, we did not observe significant mGFP lifetime changes when OBE-mGFP was co-expressed with NLS-mCherry (Figure 4L, M). Taken these observations together, we concluded that OBE3 interacts directly with SMXL5 in plant cell nuclei.

### The *OBE3* gene acts together with *SMXL3*, *SMXL4* and *SMXL5*

Since physical interaction and subcellular co-localization suggested a common action of SMXL5 and OBE3 proteins, we investigated whether the corresponding genes are functionally connected by again using root length as a first read-out for potential phloem defects. As before, *smxl4;smxl5* double mutants were short rooted, while root lengths of *smxl4* and *smxl5* single mutants were similar to wild type (Wallner et al. 2017) (Figure 5, A and B). Similarly, roots from *obe1, obe2*, *obe3* and *obe4* single mutants resembled wild type roots. In contrast, *smxl4;obe3, smxl5;obe3* and *smxl3;obe3* double mutants had short roots just as *smxl4;smxl5* (Figure 5, A-D) suggesting a concerted action of *OBE3* and *SMXL3*, *SMXL4* or *SMXL5* genes during primary root growth. Of note, we only detected a genetic interaction between *SMXL3/4/5* and *OBE3* and not between *SMXL4/5* and other *OBE* family members (Supplementary Figure 5).

**Figure 5:**
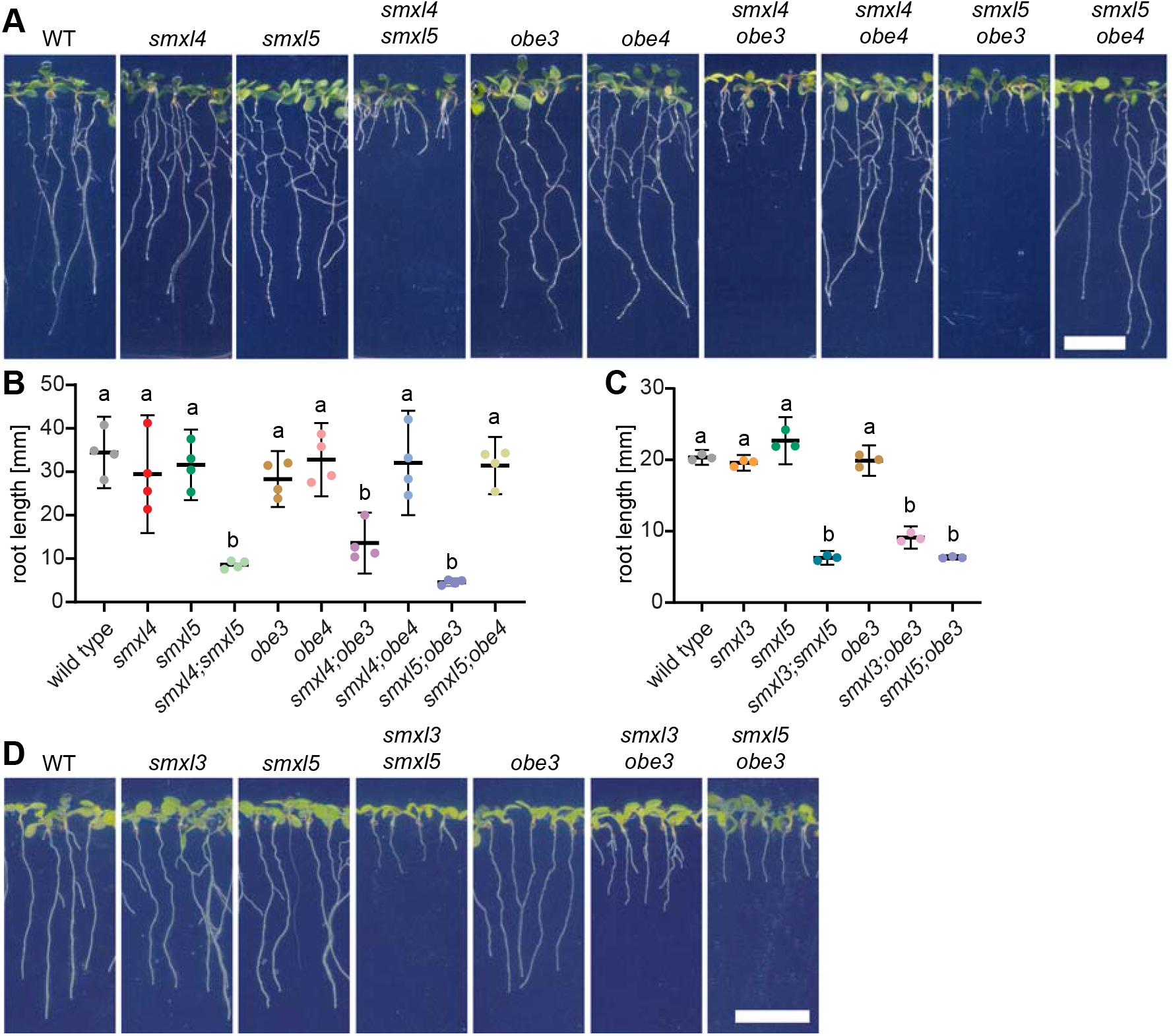
*OBE3* genetically interacts with *SMXL3/4/5* in root growth regulation. **(A)** 10 day-old wild type, *smxl4, smxl5, smxl4;smxl5, obe3, obe4, smxl4;obe3, smxl4;obe4, smxl5;obe3, smxl5;obe4* seedlings are shown from left to right. Scale bar represents 1 cm. **(B)** Quantification of root length depicted in A. Mean values of four independent experiments (n = 15-55 per experiment and genotype) were analyzed by a one-way ANOVA with post-hoc Tukey HSD (95 % CI). Statistical groups are marked by letters. **(C)** Quantification of root length depicted in D. Mean values of three independent experiments (n = 62–74 per experiment and genotype) were analyzed by a one-way ANOVA with post-hoc Tamhane-T2 (95 % CI). Statistical groups are marked by letters. **(D)** 10 day-old wild type, *smxl3, smxl5, smxl3;smxl5, obe3, smxl3;obe3, smxl5;obe3* seedlings are shown from left to right. Scale bar represents 1 cm.

### *OBE3* locally promotes early stages of phloem development

Because reduced root length of *obe3;smxl* double mutants suggested a role of *OBE3* in phloem development, we next tested whether *OBE3* is expressed in developing phloem cells by comparing the activity pattern of a translational *SMXL4:SMXL4-YFP* reporter (Wallner et al. 2017) with patterns of a translational *OBE3:OBE3-GFP* reporter (Saiga et al. 2012) (Figure 6, A-D). As reported previously (Wallner et al. 2017), the SMXL4-YFP protein accumulated specifically in nuclei of the protophloem lineage identified by enhanced Direct Red 23 staining (Figure 6A). In comparison, the *OBE3:OBE3-GFP* reporter revealed OBE3-GFP protein accumulation in nuclei of all cell types of the root tip including developing protophloem cells (Figure 6B) expressing SMXL3, SMXL4 and SMXL5 proteins (Figure 6A) (Wallner et al. 2017). We thus concluded that SMXL3/4/5 and OBE proteins had the potential to interact during early phases of phloem formation. We could not detect differences in activity patterns between *OBE3:OBE3-GFP* and *OBE4:OBE4-GFP* (Saiga et al. 2012) reporters (Figure 6, B-C) which argues against the possibility that differences in expression are the reason why *OBE3*, but not *OBE4*, genetically interacted with *SMXL3/4/5* genes.

**Figure 6:**
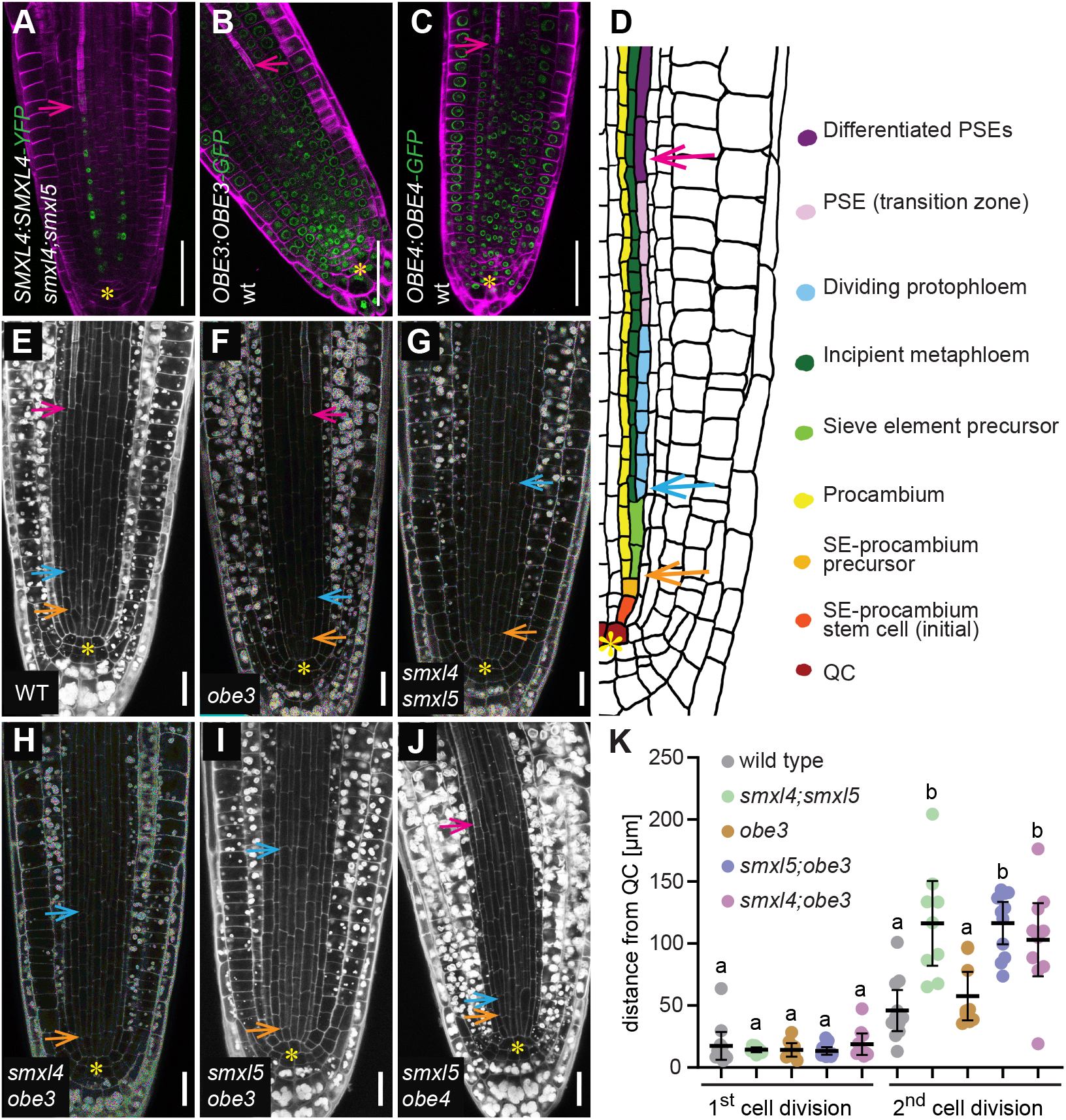
*OBE3* interacts with *SMXL5* in protophloem formation. **(A – C)** Phloem-specific activity of *SMXL4:SMXL4-YFP* (A) coincides with activity of *OBE3:OBE3-GFP* (B) and *OBE4:OBE4-GFP* (C) reporters in the developing phloem. Fluorescent signals (green) and cell wall staining by Direct Red 23 (magenta) were detected by confocal microscopy. Pink arrows point to the first differentiated SE indicated by enhanced cell wall staining. Scale bars represent 50 µm. **(D)** Schematic representation of one developing phloem pole at the root tip. Two periclinal cell divisions generate SE precursor and procambium (orange arrow) and proto- and metaphloem cell lineages (blue arrow), respectively. Differentiated SEs are marked by a pink arrow. The QC is marked by a yellow asterisk. **(E – J)** Phloem formation in 2 day-old wild type (E), *obe3* (F), *smxl4;smxl5* (G), *smxl4;obe3* (H), *smxl5;obe3* (I) and *smxl5;obe4* (J) root tips. Cell walls were stained by mPS-PI (white). Yellow asterisks mark the QC. Enhanced mPS-PI staining indicates differentiation of SEs (pink arrows) in wild type (E), *obe3* (F) and *obe4;smxl5* (G). Orange and blue arrows mark the first and second periclinal division, respectively, in the developing phloem cell lineage. Scale bars represent 20 µm. **(K)** The distance from the QC to the first and second periclinal division shown in D-I was quantified (n = 9–11). Statistical groups are indicated by letters and were determined by a one- way ANOVA with post-hoc Tukey HSD (95 % CI). Distances of 1^st^ cell divisions and 2^nd^ cell divisions were compared independently.

To evaluate whether growth defects observed in *smxl;obe3* roots are correlated with the same type of protophloem defects observed in *smxl4;smxl5* mutants, we analyzed phloem development in the respective genetic backgrounds. During protophloem development, SE procambium-precursors divide periclinally to give rise to procambium and SE precursor cells. After 2-3 anticlinal divisions, SE precursor cells divide again periclinally to initiate meta- and protophloem cell files that subsequently undergo gradual differentiation (Figure 6D) (Wallner et al. 2017). Our analysis revealed that in *obe3* mutant both the periclinal cell divisions and the onset of SE differentiation appeared as in wild type roots (Figure 6, E, F, K). In contrast, in *smxl4;obe3* and *smxl5;obe3* mutants, the onset of the second periclinal division initiating meta- and protophloem cell lineages was similarly delayed as in *smxl4;smxl5* mutants and the enhanced mPS-PI staining visualizing differentiated SEs was likewise absent (Figure 6 G-I, K and Supplementary Figure 6). This observation demonstrated that, like the more locally expressed *SMXL3/4/5* genes, *OBE3* substantially contributes to phloem formation. Moreover, because respective single mutants did not show these defects (Figure 6 and Supplementary Figure 6), we concluded that *OBE3* or *SMXL5*-deficient plants represent sensitized backgrounds for the functional loss of the other regulator. In accordance with the wild type-like root growth of those plants, protophloem formation in *smxl5;obe4* double was indistinguishable from wild type (Figure 6, E and J).

Because *OBE3* is broadly expressed, phloem defects observed in *smxl5;obe3* mutants could arise due to a function of *OBE3* in other tissues than developing phloem cells, which would contradict a direct interaction of SMXL5 and OBE3 proteins. To address this concern and to determine whether *OBE3* acts cell-autonomously on phloem development, we expressed *OBE3* exclusively in developing phloem cells by introducing a transgene driving an OBE3-turquoise fusion protein under the control of the *SMXL5* promoter (*SMXL5:OBE3- turquoise*) into a *smxl5;obe3* double mutant background. Microscopic analysis of root tips from *smxl5;obe3/SMXL5:OBE3-turquoise* lines confirmed the presence of the OBE3-turquoise protein in nuclei of developing protophloem cells (Figure 7, A and B) as described for the SMXL5 protein expressed under the control of the same promoter (Wallner et al. 2017). When comparing *smxl5;obe3/SMXL5:OBE3-turquoise* lines with *smxl5;obe3* mutants, we observed that expression of *OBE3-turquoise* within the *SMXL5* domain was indeed sufficient to restore root length in *smxl5;obe3* double mutants (Figure 7, C and D). Additionally, enhanced Direct Red staining of the mature protophloem indicating SE differentiation was recovered in *smxl5;obe3* carrying the *SMXL5:OBE3-turquoise* transgene (Figure 7, A and B). To see whether the predominant reduction of *OBE3* activity in developing phloem cells is furthermore sufficient for generating phloem defects, we designed two artificial microRNAs targeting the *OBE3* mRNA (*obe3-miRNAs*) (Schwab et al. 2006) and expressed them independently under the control of the *SMXL5* promoter in *smxl5* mutant plants. As expected, the majority of those plants was short rooted, indicating that *OBE3* knock-down within the *SMXL5* domain was sufficient to evoke a *smxl5;obe3*-like phenotype in *smxl5* mutants (Figure 7, C and D). We thus concluded that *OBE3* fulfils a cell autonomous and *SMXL5*-dependent role in protophloem formation.

**Figure 7:**
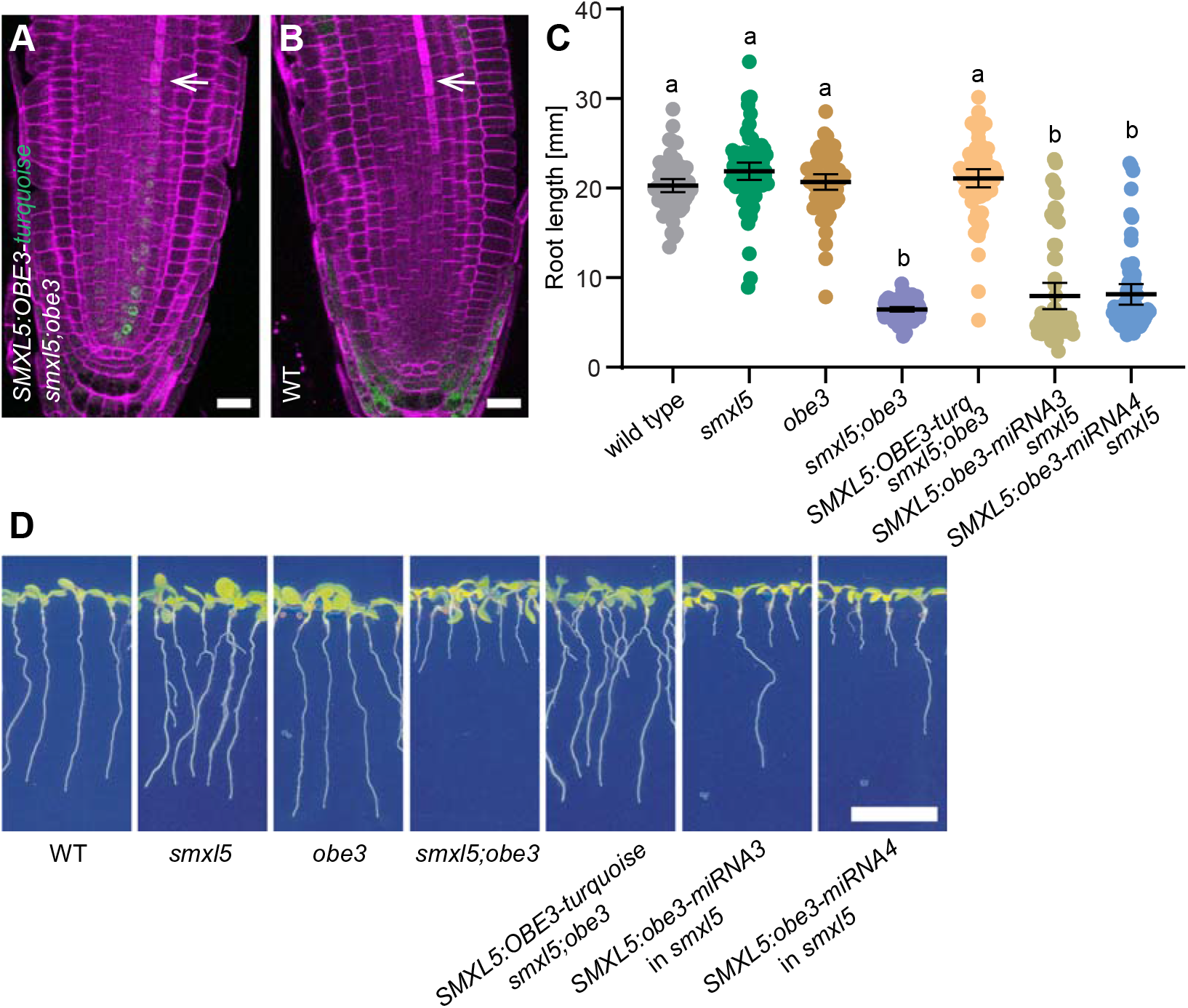
Phloem-specific *OBE3* expression is sufficient for promoting root growth. **(A – B)** Seven day-old *smxl5;obe3* root tips carrying an *SMXL5:OBE3-turquoise* reporter (green signal, A) were compared to a non-transformed wild type (WT) root tip (B). Cell walls were stained by DirectRed (magenta). White arrows mark differentiated SEs. Scale bars indicate 20 µm. **(C)** Root length quantification of plants depicted in D. Results from one representative experiment out of three independent experiments (n = 61–71 per experiment and genotype) are shown. Mean values were analyzed by one-way ANOVA with post-hoc Tamhane-T2 (95 % CI). Statistical groups are indicated by letters. **(D)** 10 day-old wild type, *smxl5, obe3, smxl5;obe3, SMXL5:OBE3-turquoise*/*smxl5;obe3*, *SMXL5:obe3-miRNA3*/*smxl5* and *SMXL5:obe3-miRNA4*/*smxl5* seedlings are shown from left to right. Scale bar represents 1 cm.

### *SMXL* and *OBE3* genes determine chromatin structure in phloem cells

To probe the common role of SMXL and OBE3 proteins and their putative effect on chromatin signatures (Saiga et al. 2012; Ma et al. 2017; Wang et al. 2020), we performed phloem-specific Assays for Transposase-Accessible Chromatin using sequencing (ATAC-seq) revealing chromatin structure in a genome-wide fashion (Buenrostro et al. 2015). To this end, we fluorescently labeled nuclei in phloem-associated cells in wild type, *smxl5*, *smxl4;smxl5* and *smxl5;obe3* plants by expressing a histone H4-GFP protein fusion under the control of the *SMXL5* promoter (*SMXL5:H4-GFP*, (Shi et al. 2019), Supplementary Figure 7A). Isolation of nuclei from those lines and their fluorescence-activated sorting into GFP-positive and GFP- negative populations (FANS (Shi et al. 2021), Supplementary Figure 7B) allowed ATAC-seq analyses separately on phloem-related and non-phloem-related cells. Thereby, we detected 36,925 to 40,168 open chromatin regions (OCRs) in all sample types mostly located in 5’ and 3’ regions of respective gene bodies (Figure 8A, B, Supplementary Figures 8, 9 and 10). Importantly, the overall conformation was similar in all samples (Supplementary Figure 8) suggesting, by large, a comparable chromatin structure in phloem and non-phloem cells and an independency of most chromatin domains from *SMXL4*, *SMXL5* or *OBE3* activity. Among 38,562 OCRs detected in phloem cells from wild type, 1488 OCRs associated with 1072 genes showed a significant increase of read alignment in comparison to non-phloem cells (Figure 8A; Supplementary Dataset 1 and 2; fold change > 2, poisson enrichment *p*-value < 0,05, using 40 M reads). These genes included OCRs in promoter regions of the phloem-specific *OPS*, *BAM3*, *CVP2*, and *APL* genes (Supplementary Figures 9 and 10, Supplementary Dataset 1 and 2) suggesting that we succeeded in phloem-specific ATAC-seq analysis. In accordance with the wild type-like phenotype of *smxl5* mutants, 1055 of 36,925 OCRs associated with 809 genes showed a significant increase of read alignment in the *smxl5* phloem in comparison to non-phloem cells in *smxl5* again including *OPS*, *BAM3*, *CVP2*, and *APL* (Figure 8A, Supplementary Figures 9 and 10, Supplementary Datasets 1 and 2). In contrast, only 48 and 316 phloem-associated OCRs were detected in *smxl4;smxl5* and *smxl5;obe3* double mutants, respectively (Figure 8A, Supplementary Datasets 1 and 2), suggesting a loss of phloem-specific chromatin signatures in both backgrounds.

**Figure 8:**
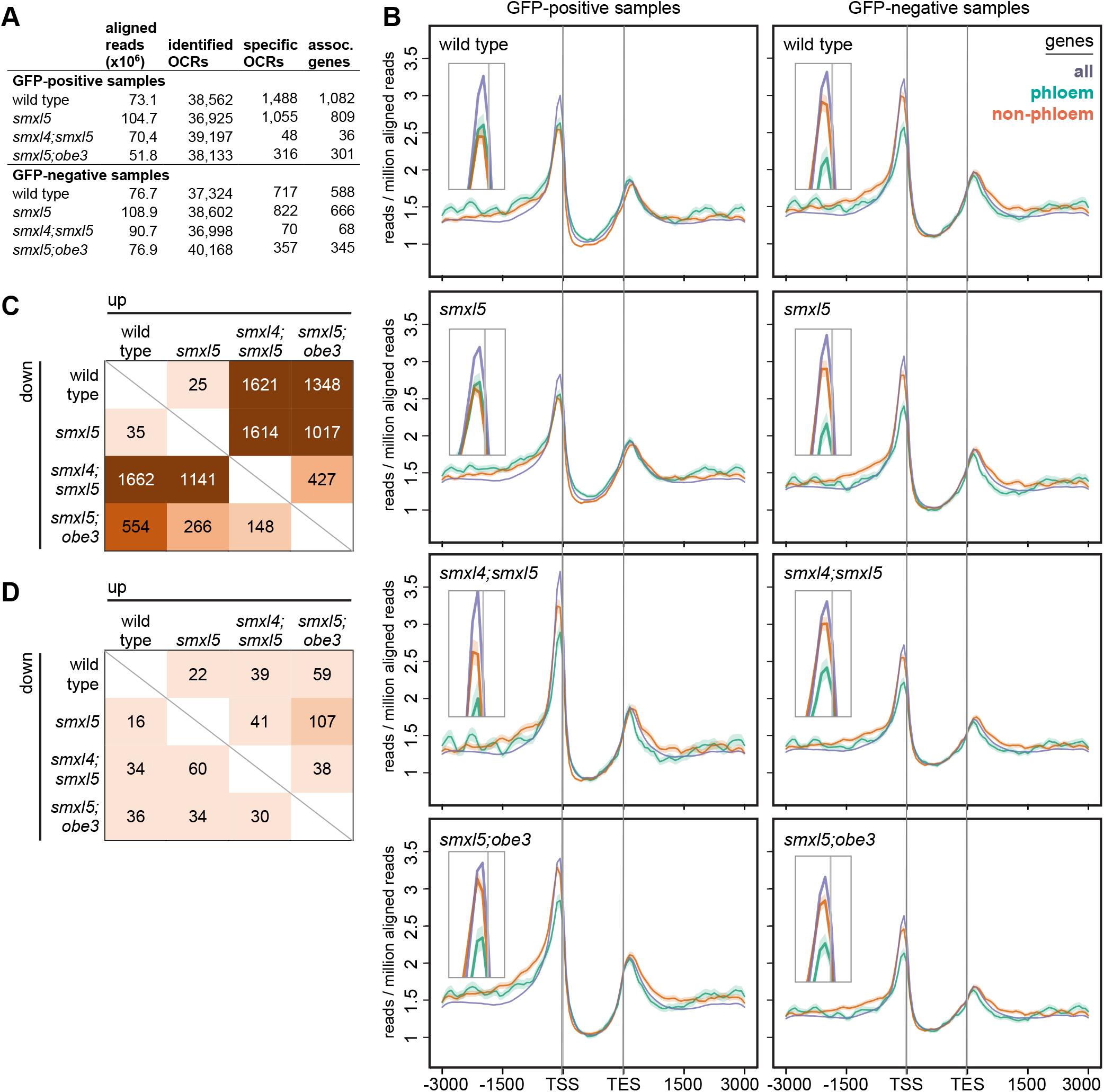
*SMXL* and *OBE3* genes determine phloem-related chromatin profile. **(A)** Summary of ATAC-seq results for the different samples. Specific OCRs were identified by comparing GFP-positive and GFP-negative samples of the respective genetic backgrounds. After initial alignment, OCR detection and other subsequent analyses were conducted using 40 M reads aligning to the nuclear genome excluding duplicates for each sample. **(B)** Chromatin conformation profiles for all genes found in the Arabidopsis genome or only for phloem and non-phloem genes according to Brady et al., 2007. Read alignment was adjusted to transcriptional start sites (TSS) and transcriptional end sites (TES). X-axes show distance to TSSs and TES in base-pairs. Inserts in all graphs are enlargements of the peaks identified close to TSSs. **(C, D)** Comparison of OCR profiles found in GFP-positive (C) and GFP-negative (D) samples. Numbers indicate the number of OCRs with significantly more (‘up’) or less (‘down’) aligned reads.

This conclusion was also supported by direct comparison of the OCR profile obtained from GFP-positive samples. In total, only 60 OCRs showed significantly more or fewer aligned reads comparing *smxl5* mutants with wild type (Figure 8C, Supplementary Dataset 3, fold change >2, poisson enrichment *p*-value < 0.05, using 40 M reads) arguing for a high similarity of the chromatin profile in phloem-related cells from both backgrounds. In contrast, *smxl4;smxl5* and *smxl5;obe3* double mutants differed considerably from both wild type and *smxl5* mutants with regard to their phloem-related chromatin profile. *smxl4;smxl5* mutants showed 3283 OCRs in total with significant differences in read alignment in comparison to wild type and, for *smxl5;obe3* mutants, 1902 differential OCRs were found. A similar situation was found for both double mutants in comparison to *smxl5* mutants (Figure 8C, Supplementary Dataset 3). Importantly, the number of differential OCRs was lower when comparing *smxl4;smxl5* and *smxl5;obe3* samples directly. In this case, only 575 differential OCRs were detected (Figure 8C) suggesting that both double mutants differed similarly from wild type and *smxl5* mutants with regard to their chromatin profile in phloem-related cells. Beyond the overall OCR profile, the chromatin around phloem-related genes like *OPS*, *BAM3*, *CVP2*, and *APL* displayed a more condensed conformation in GFP-positive nuclei from *smxl4;smxl5* and *smxl5;obe3* and the difference between GFP-positive and -negative nuclei was less pronounced in these cases (Supplementary Figure 9 and 10).

When comparing the OCR profile obtained from GFP-negative samples, the difference between the genetic backgrounds was substantially lower with a maximum of 141 differential OCRs comparing *smxl5* and *smxl5;obe3* backgrounds (Figure 8D, Supplementary Dataset 4). Together, these comparisons suggested a considerable and mostly phloem-specific difference between chromatin conformation in phloem-related cells when comparing wild type and *smxl5* mutants on the one side and *smxl4;smxl5* and *smxl5;obe3* double mutants on the other side. Moreover, we concluded that phenotypic changes are similar in *smxl4;smxl5* and *smxl5;obe3* mutants, underlining the functional relatedness of *SMXL* and *OBE3* genes.

To see how chromatin conformation differed overall between the different genotypes, we defined a group of ‘phloem’ genes (combined SUC2, APL and S32 domain genes according to (Brady et al. 2007), Supplementary Dataset 5) and ‘non-phloem’ genes (combined M1000, cortex, COBL9, GL2, AGL42, PET111, and LRC domain genes according to (Brady et al. 2007), Supplementary Dataset 5). When analyzing chromatin conformation associated with the different gene groups, a similar pattern in wild type and *smxl5* mutants was observed with a more open chromatin around transcriptional start sites (TSS) of non-phloem genes in GFP- negative samples compared to GFP-positive samples (Figure 8B). Interestingly, in GFP- positive samples from *smxl4;smxl5* and *smxl5;obe3* double mutants, overall chromatin conformation of phloem genes was similar to wild type. In contrast, chromatin of non-phloem genes was more open around TSSs in GFP-positive samples and the conformation resembled the pattern in GFP-positive samples (Figure 8B). This observation again indicated that phloem-specific chromatin signatures were equally reduced in phloem-related cells of *smxl4;smxl5* and *smxl5;obe3* double mutants reflecting a comparable function of both SMXL4/5 and OBE3 proteins.

## Discussion

Cell type specification is fundamental for establishing multicellular organisms and, in recent years, the phloem has become an instructive model for studying this aspect in plants (Anne and Hardtke 2017; Blob et al. 2018; Gujas et al. 2020; Marhava et al. 2020). With our study we provide new insights into the regulation of (proto)phloem formation by revealing a role of the putative chromatin remodeling protein OBE3 (Saiga et al. 2012) and a direct interaction between the OBE3 protein and the central phloem regulator SMXL5. Based on our findings, we propose that SMXL proteins fulfil their role in phloem formation by creating a distinct chromatin signature important for establishing phloem identity.

### SMXL3/4/5 and OBE3 proteins act together on early stages of phloem development

SMXL3/4/5 and OBE3 proteins are already expressed in phloem stem cells (Wallner et al. 2017), which was so far not described for other phloem regulators (Blob et al. 2018). Here, we detected OPS and BRX protein accumulation already in those stem cells raising the question of functional interdependence. The positive effect of *SMXL4* and *SMXL5* gene functions on OPS and BRX protein accumulation in those and more mature phloem cells suggests that *SMXL* genes are required for the establishment of a phloem-specific developmental program including *OPS* and *BRX* gene activities. The different subcellular localization of OPS, BRX and BAM3 proteins on the one side and SMXL proteins on the other side argues against a more interconnected mechanism of both groups of regulators. The conclusion that both groups act on different aspects of phloem formation is furthermore supported by our genetic analyses which revealed a combination of distinct phloem defects in respective double mutants. Here, we propose that SMXL proteins fulfil their role in the nucleus of phloem stem cells and beyond by direct interaction with OBE3. In fact, the *SMXL3/4/5-OBE3* interaction seems to be a prerequisite for both the initiation of phloem cell fate and a timely onset of differentiation. On the genetic level, *smxl4;obe3*, *smxl5;obe3* and the *smxl4;smxl5* double mutants share the same phloem defects meaning that they are deprived of protophloem formation within the RAM. Other reported phloem mutants have either problems with completing phloem differentiation in general, which is the case in mutants of the *ALTERED PHLOEM DEVELOPMENT* (*APL*) gene (Bonke et al. 2003; Truernit et al. 2008; Kondo et al. 2016), or they develop ‘gap cells’ in which SE-differentiation is disturbed as in *ops* or *brx* single or in *cvp2;cvp2-like1* (*cvl1*) double mutants (Depuydt et al. 2013; Anne et al. 2015; Rodriguez-Villalon et al. 2015; Marhava et al. 2018; Breda et al. 2019). In contrast, *smxl5;obe3*, *smxl4;obe3* and multiple mutants of the *SMXL3/4/5* genes show absence of all morphological hallmarks of phloem formation within the RAM. This suggests that *SMXL3/4/5* and *OBE3* act together on the establishment of a phloem-specific developmental program. Interestingly, complete suppression of protophloem formation was so far only reported for roots treated with certain CLAVATA3/ESP-RELATED (CLE) peptides, such as CLE45, which signal through the leucine-rich repeat receptor like kinase BAM3 and the pseudokinase CORYNE (CRN) (Depuydt et al. 2013; Hazak et al. 2017). However, the formation of ‘gap cells‘ in *ops* or *brx* mutants is suppressed in *BAM3-*deficient backgrounds (Rodriguez-Villalon et al. 2014). This shows that none of those factors is required to obtain protophloem cell identity and proper differentiation in the first place (Depuydt et al. 2013; Rodriguez-Villalon et al. 2014). Moreover, the epistatic relationship between *SMXL4/5* and *BAM3* genes as observed in this study shows that the SMXL and BAM3 pathways are functionally independent.

### A putative role of a SMXL3/4/5/OBE3 protein complex in chromatin remodeling

PHD-finger motifs as carried by the OBE3 protein are known to be epigenetic readers binding to histone H3 tails carrying distinct post-translational modifications such as trimethylation of lysine 4 (H3K4me3) or lysine 9 (H3K9me) marking actively transcribed or silent chromatin regions, respectively (Sanchez and Zhou 2011). Although PHD finger proteins themselves are not necessarily activating or repressing, they can indirectly modify transcription by recruiting chromatin modifying complexes (De Lucia et al. 2008). Indeed, OBE proteins have been proposed to remodel chromatin structure during embryogenesis and thereby transcriptionally activate RAM initiation factors (Saiga et al. 2012). SMXL proteins share an ethylene-responsive element binding factor-associated amphiphilic repression (EAR) motif which interacts with transcriptional regulators of the TOPLESS (TPL) family (Pauwels et al. 2010) and they have recently shown to bind DNA directly (Wang et al. 2020). Indeed, the strigolactone signaling mediators SMXL6, SMXL7, SMXL8 and DWARF53 (D53), an SMXL protein from rice, directly interact with TPL-like proteins (Soundappan et al. 2015; Wang et al. 2015; Ma et al. 2017). TPLs are transcriptional co-repressor which can recruit histone deacetylases (HDAC) and, thereby, induce chromatin condensation and transcriptional suppression (Krogan and Long 2009; Ma et al. 2017). In addition to EAR motifs, SMXLs share a conserved double caseinolytic protease (Clp) domain with ATPase activity that resembles heat shock protein 101 (HSP101) (Jiang et al. 2013; Zhou et al. 2013). As recently proposed, the p-loop ATPase domain of D53 fosters the formation of TPL hexamers and the threading of DNA through a central pore of this hexamer inducing nucleosome repositioning and/or higher-order chromatin reorganization (Ma et al. 2017).

Judging from the protein domains, their sub-nuclear localization and the similar loss of phloem-specific chromatin signatures in *smxl4;smxl5* and *smxl5;obe3* mutants, we hypothesize that SMXL3/4/5 and OBE3 form a protein complex that is involved in chromatin remodeling and/or transcriptional regulation of downstream targets. In summary, our study reveals the functional interaction between *SMXL3/4/5* and *OBE3*, an important step in understanding plant tissue formation.

## Material and methods

### Plant material

Genotypes of plant species *Arabidopsis thaliana* (L.) Heynh. of the ecotype Columbia (Col) used for genetic analysis are listed in the Key Resource Table. Sterile seeds were stratified in microcentrifuge tubes containing dH2O at 4 °C in the dark for 3 days and then sown in rows on ½ Murashige and Skoog (MS) medium-plates supplemented with 1 % sucrose and grown vertically. Seedlings were grown in long day (LD, 16 h light and 8 h dark) conditions at 21 °C for 2-10 days. Flowering plants were transformed by the floral dip method described earlier (Clough and Bent 1998). Seeds were liquid sterilized by 70 % ethanol supplemented with 0.2 % Tween-20 for 15 min, washed twice with 100 % ethanol and air dried under sterile conditions. *bam3-2* (SALK_044433), *obe1-1* (SALK_075710), *obe3-2* (SALK_042597) and *obe4-1* (SALK_082338) mutants were obtained from the NASC stock center. *smxl3-1* (SALK_024706), *smxl4-1* (SALK_037136), *smxl5-1* (SALK_018522), and *obe2-2* mutants were described before (Thomas et al. 2009; Wallner et al. 2017). *smxl4-1;smxl5-1;bam3-3* triple mutants were generated by CRISPR/Cas9 targeting *BAM3* in a *smxl4-1;smxl5-1* mutant background (creating the *bam3-3* allele), leading to a G insertion after position 707 of the CDS and to the generation of a stop codon after 255 aa of the BAM3 protein (i.e. in the LRR receptor domain). All other lines are referenced in the text.

### Yeast-Two-Hybrid

The yeast-based screen for proteins interacting with SMXL5 was performed by Hybrigenics (Evry, France) as described before (Legrain et al. 2001). The yeast strain *AH109* was used for the yeast two-hybrid assay according to Matchmaker^TM^ Two-Hybrid System 3 (Clontech, Palo Alto) and grown on YPD (full medium) or SD (selective medium)-agar plates for 3-5 days at 28°C, then stored at 4°C and stroked onto new plates every 10 days. Dilution series (OD600 1-0.001) of transformed yeast strains were grown for 3 days on selective medium (SD) -Leu/-Trp/-His/-Ade selecting for protein interaction or SD -Leu/-Trp selecting for the presence of the plasmids. When grown in liquid YPD medium for transformation, yeast was grown over night at 28 °C with shaking at 250 rpm. For expressing the GAL4BD-SMXL5 fusion protein in yeast the open reading frame of the *SMXL5* gene was cloned into XmaI/BamHI sites of the *pGBKT7* plasmid (Clontech, Palo Alto) resulting in *pEW6*. For expressing the GAL4AD- OBE3 fusion protein, the open reading frame of the *OBE3* gene was cloned into XmaI/BamHI sites of the *pGADT7* plasmid (Clontech, Palo Alto) resulting into *pEW11*.

### Agrobacterium tumefaciens

The *Agrobacterium tumefaciens* genotypes C58C1: RifR with pSoup plasmid (TetR) or ASE: KanR, CamR with pSoup+ plasmid (TetR) were used for transformation of *Arabidopsis thaliana* or infiltration of *N. benthamiana* leaves and grown at 28 °C over night in liquid YEB medium on a shaker (180 rpm to an OD600 > 1) or plated on YEB-plates and grown in an incubator (Fraley et al. 1983; Ashby et al. 1988; Hellens et al. 2000). Antibiotics were used for plasmid selection.

### Root length measurements

For measuring root lengths, seedlings were scanned by a commercial scanner and analyzed using ImageJ 1.49d (Schindelin et al. 2012). For CLE45 treatments, plants were germinated on normal MS media (mock) or media containing 50 nM CLE45 and root length measurements were performed after five days.

### Genotyping

Genotyping was performed by PCR using primers listed in Supplementary Table 1. Further information about standard DNA extraction and genotyping can be found in (Wallner 2018).

### Transient protein expression in *N. benthamiana*

*N. benthamiana* plants were used for transient protein expression and grown in the greenhouse at approximately 25 °C and watered daily. Transformed *Agrobacteria* were stored as glycerol stocks and grown in a 10 ml YEB liquid culture prior to use. The densely grown culture was centrifuged at 4000 rpm for 5 min at RT. The supernatant was removed and the pellet was washed with 5 ml induction buffer and re-suspended in 10 ml induction buffer. Culture densities were adjusted to an OD600 of 1. Prior to infiltration, these bacterial solutions were mixed with *Agrobacteria* expressing *35S:P19* in a ratio 1:2 and incubated in the dark for 2-3 h (Voinnet et al. 2003; Scholthof 2006). *N. benthamiana* leaves were infiltrated with the mixtures using a 1 ml syringe (Becton Dickinson S.A., Heidelberg, Germany). Leaves were harvested three days after infiltration.

### Protein extraction, immunoprecipitation and Western blot

Infiltrated *N. benthamiana* leaves were frozen in liquid nitrogen and ground by a mortar. Proteins were extracted by mixing the leaf powder 1:1 with extraction buffer (50 mM Na3PO4, 150 mM NaCl, 10% glycerol, 5 mM EDTA, 10 mM β-mercaptoethanol, 0.1% triton X-100, 2 mM NaVO4, 2 mM NaF, 20 µM MG-132, 1 mM PMSF, 1x cOmpleteTM Protease Inhibitor Cocktail (Roche; Basel, Switzerland)). Each sample was vortexed for 10 sec and centrifuged at 13000 rpm for 10 min at 4 °C. The protein extract was retrieved by sieving it through a nylon mesh. Protein quantities were measured by Bradford assays according to the manual provided with the Bio-Rad Protein Assay Dye Reagent Concentrate (Bio-Rad Laboratories; Hercules, USA). Proteins were immunoprecipitated by 50 µl Anti-HA MicroBeads (Miltenyi Biotec, Bergisch Gladbach, Germany) after incubation for 2.5 h at 4 °C while slowly rotating. Beads were captured by µ Columns (Miltenyi Biotec, Bergisch Gladbach, Germany) on magnetic stands by following the user manual and washed three times by 200 µl Wash buffer I (extraction buffer without β-mercaptoethanol) and two times by 200 µl Wash buffer II (50 mM Na3PO4, 150 mM NaCl, 10% glycerol, 5 mM EDTA). Proteins were eluted by 2x Laemmli buffer (95 °C) and separated by size on a SDS-PAGE with subsequent western blotting. Detailed procedures can be found in (Wallner 2018). SMXL5-3xHA and 6xMyc-OBE3 bands were detected by antibodies Anti-HA-Peroxidase High Affinity (3F10) (Roche; Basel, Switzerland) or c-Myc Antibody (9E10) sc-40 HRP (Santa Cruz Biotechnology, Santa Cruz, USA), respectively and visualized by chemiluminescence agents SuperSignal™ West Femto Maximum Sensitivity Substrate (Thermo-Scientific; Waltham, USA) by an Advanced Fluorescence and ECL Imager (Intas Science Imaging Instruments, Göttingen, Germany).

### FRET-FLIM analyses

FRET-FLIM analyses were performed principally as described previously (Ladwig et al. 2015; Arongaus et al. 2018). Briefly, measurements were performed using a Leica TCS SP8 microscope (Leica Microsystems, Germany) equipped with a rapidFLIM unit (PicoQuant). Images were acquired using a 63x/1.20 water immersion objective. For the excitation and emission of fluorescent proteins, the following settings were used: mGFP at excitation 488 nm and emission 500-550 nm; and mCherry at excitation 561 nm and emission 600-650 nm. The lifetime τ [ns] of either the donor only expressing cells or the cells expressing the indicated combinations was measured with a pulsed laser at an excitation light source of 470 nm and a repetition rate of 40 MHz (PicoQuant Sepia Multichannel Picosecond Diode Laser, PicoQuant Timeharp 260 TCSPC Module and Picosecond Event Timer). The acquisition was performed until 500 photons in the brightest pixel were reached. To obtain the GFP fluorescence lifetime, data processing was performed with SymPhoTime software and bi-exponential curve fitting and a correction for the instrument response function. Statistical analysis was carried out using the JMP 14 software (JMP, USA).

### Direct Red 23 staining

To preserve fluorescent signals in roots, seedlings were fixed in a vacuum chamber for 1 h by 4 % (w/v) PFA dissolved in PBS. The tissue was washed twice by PBS and cleared with ClearSee solution for a minimum of two days according to (Kurihara et al. 2015). Cleared seedlings were stained by 0.01 % (w/v) Direct Red 23 in ClearSee solution for 1 h. Excess staining was removed by clearing once again in pure ClearSee solution for 1 h.

### mPS-PI staining

The mPS-PI staining of roots was carried out as described before (Truernit et al. 2008).

### Confocal microscopy

For confocal microscopy, TCS SP5 or SP8 microscopes (Leica Microsystems; Mannheim, Germany) were used. GFP signals were excited at excited by an argon laser at 488 nm, collecting the emission between 500-575 nm. YFP was excited by an argon laser at 514 nm and the emission detected in a range of 520-540 nm. DirectRed stained tissue was excited at 561 nm (DPSS laser) and emission was detected at wavelengths >660 nm. mPS-PI stained tissue was excited at 561 nm (DPSS laser) and emission was detected at 590-690 nm.

### Molecular cloning and miRNA generation

*OPS:SMXL5-VENUS* (*pNT52*)*, OPS:ER-VENUS* (*pNT53*), *BAM3:SMXL5-VENUS* (*pNT49*)*, BAM3:ER-VENUS* (*pNT50*)*, CVP2:SMXL5-VENUS* (*pNT16*)*, CVP2:ER-VENUS* (*pNT69*)*, APL:SMXL5-VENUS* (*pNT10), APL:ER-VENUS (pNT68), SMXL4:BRX-VENUS* (*pNT72*), *SMXL5:OBE3-turquoise* (*pEW72*)*, 35S:5xc-Myc-OBE3* (*pEW78*)*, 35S:SMXL5- mCherry* (*pVL122*)*, 35S:OBE3-mGFP* (*pVL127*)*, 35S:mCherry-NLS* (*pMG103*) and *UBI10:mGFP-mCherry-NLS* (*pCW194*) constructs were generated by using appropriate modules according to the GreenGate manual described in (Lampropoulos et al. 2013). Destination modules, entry modules, and correlating primers for amplifying DNA fragments for generating entry modules are depicted in Supplementary Table 1. In case reporter proteins were targeted to the endoplasmatic reticulum (ER), they were fused to the appropriate motifs (Haseloff et al. 1997). *miRNAs* targeting *OBE3* transcripts were designed and cloned according to the manual provided by the WMD3 - Web MicroRNA Designer Version 3 (Max Planck Institute for Developmental Biology, Tübingen. http://www.weigelworld.org) with primers listed in Supplementary Table 1. To generate *35S:SMXL5-3xHA* (*pEW33*), the *SMXL5* CDS was amplified by primers listed In Supplementary Table 1 and cloned into BamHI/XbaI sites of *pGreen0229:35S* (Hellens et al. 2000) resulting in *pKG33*. Next, ssDNA sequences coding for 3xHA (Supplementary Table 1) were annealed by gradual cool-down from 80 °C to 50 °C and inserted into vector *pKG33* using BamHI/XmaI sites, resulting in plasmid *pEW31*. Further information about detailed cloning procedures can be found in (Wallner 2018).

### Fluorescence Activated Nucleus Sorting (FANS) and ATAC-seq

Nucleus extraction for FANS/ATAC-seq was based on (Thibivilliers et al. 2020) and carried out on ice. Approximately 1 gr of three week-old Arabidopsis seedlings grown in ½ MS LD conditions were collected in a 60 mm petri dish and thoroughly chopped using a razor blade for five minutes in nucleus isolation buffer (1x NIB, CelLytic™-PN Isolation/Extraction buffer, Sigma-Aldrich, cat.no. CELLYTPN1) supplemented with 10 µg/ml Hoechst 33342 (Sigma-Aldrich, B2261) as the final concentration. After incubation for 15 min at 4°C in darkness, the nucleus suspension was applied to 50 µm nylon strainers mounted into a 30 µm nylon strainers at the top (Sysmex, CellTrics), and filtered further by a FACS tube with 35 µm strainer cap (Corning, #352235). 15,000 nuclei for each sample were then sorted according to their GFP signal levels into a 300 µL collection buffer (15 mM TRIS-HCl ph 7.5, 20 mM NaCl, 80 mM KCl, 0.1 % Triton) by a BD FACSAriaTM IIIu cell sorter using a 100 µm sort nozzle and a 30 kHz drop drive frequency. The gate for GFP+ nuclei was set using wild type as a reference (Supplementary Figure 7B). Next, samples were centrifuged at 3000 g for 10 minutes at 4°C. The supernatant was partially removed and the nuclei resuspended in 300 µL Tris-Mg buffer (Tris HCl pH8 10 mM, MgCl2 5 mM). A second washing step was performed using centrifugation at 1000 g for 10 minutes at 4°C and the supernatant was removed completely. For tagmentation, the TDE1 (Nextera Tn5 Transposase) Tagment DNA Enzyme (Illumina kit #20034198) was immediately applied to the samples by resuspending the nuclei in the transposition reaction mix and the suspension was incubated for 30 minutes at 37°C while gently shaking (Buenrostro et al. 2015). The samples were then purified (NEB, Monarch Nucleic Acid Purification Kit #20034198), and 12 µl of the transposed DNA were used for amplification by PCR (8 µl H2O, 2.5 µl 25 µM Customr Nextera PCR primer 1 (Buenrostro et al. 2013), 2.5 µl 25 µM Custom Nextera PCR Primer 2 and 25 µl NEB Next High Fidelity 2x PCR Master Mix (NEB, #M0541) with 1 cycle of (72°C for 5 min, 98°C for 30 sec) and 16 cycles of (98°C for 10 sec, 63°C for 30 sec, 72°C for 1 min). The library was cleaned up using the Coulter Agencourt AMPure XP kit (Beckman-Coulter, #10136224) applying a 1:1 vol:vol (PCR product:beads) and eluted in 15 µl 15 mM Tris. The library was sequenced using NextSeq 550 (Illumina) using High-Output with 40 cycles Paired End mode.

### Analysis of sequencing data

Reads were mapped to Arabidopsis TAIR 10 genome by bowtie2 (v2.2.6) (Langmead and Salzberg 2012). Output sam files were converted to bam files and then trimed with “-q 2” and “1 2 3 4 5” parameters in samtools (v0.1.19) (Li et al. 2009) to remove mitochondrial and chloroplast sequence. Output bam files were treated with picard MarkDuplicates (v2.25.5) to remove duplicated reads. Then, output files were passed through samtools view again, to obtain 40 M reads (targeted) for further analysis. For the profiles, bigWig files were generated by bamCoverage in deepTools (v3.5.1) (Ramírez et al. 2014) from bam files with 10 bp bin size, and visualized by igv (Thorvaldsdóttir et al. 2013). For peak calling, bam files were converted to TagDirectory by homer (v4.10.3) (Heinz et al. 2010), and peaks were called by findPeaks in homer with “-region -minDist 150” option. Differential peaks were obtained by getDifferentialPeaks function with “-F 2.0 -P 0.05” option. TSS profiles were made by ngs.plot (v2.47) (Shen et al. 2014). Raw data were deposited to the NCBI GEO archive (Barrett et al. 2013) under the accession GSE184344 (https://www.ncbi.nlm.nih.gov/geo/query/acc.cgi?acc=GSE184344).

### Quantification and statistical analyses

Statistical analyses were performed using IBM SPSS Statistics for Windows, Version 22.0. Armonk, NY: IBM Corp or using GraphPad Prism version 6.01 (GraphPad Software, La Jolla, USA). Means were calculated from measurements with sample sizes as indicated in the respective figure legends. In general, all displayed data represents at least three independent, technical repetitions, unlike otherwise indicated. Error bars represent ± standard deviation. All analyzed datasets were prior tested for homogeneity of variances by the Levene statistic. One-way ANOVA was performed, using a confidence interval (CI) of 95% and a post-hoc Tukey HSD for comparisons of five or more data sets of homogenous variances or a post-hoc Tamhane-T2 in case variances were not homogenous. Graphs were generated in GraphPad Prism version 6.01 or 9.2 (GraphPad Software, La Jolla, USA) or in Excel (Microsoft, Redmond, USA).

## Acknowledgements

This work was supported by the SFB 1101 to K.H., J.U.L, T.G. and SFB 873 (Deutsche Forschungsgemeinschaft, DFG) to J.U.L. and T.G., a Heisenberg Professorship (DFG, GR 2104/5-2), and an ERC Consolidator grant (PLANTSTEMS, #647148) to T.G. , a postdoctoral fellowship of the Japan Society for the Promotion of Science [JSPS Overseas Research Fellowships 201960008 to D.S.], and a PhD student fellowship of the Cusanuswerk to N.T.. We are grateful to Christian Hardtke (University of Lausanne, Switzerland), Dolf Weijers, Shunsuke Saiga (both at Wageningen University, The Netherlands), Andy Maule (John Innes Centre, UK), Sebastain Wolf and Karin Schumacher (both Heidelberg University, Germany) for providing seed material and constructs. We are also grateful to Monika Langlotz (Cell Networks Flow Cytometry & FACS Core Facility, Heidelberg University, Germany) and David Ibberson (Cell Networks Deep Sequencing Core Facility, Heidelberg University, Germany) for providing excellent technical support.

## Author Contributions

Conceived and designed the experiments: EW, NT, FW, VL, DS, LL, KH, TG. Performed experiments: EW, NT, FW, LL, IJ, VL, PH, YX, MG, CW. Analysed the data: EW, NT, FW, KH, JUL, DS, TG. Wrote the paper: EW, TG.

## Supplementary item titles and legends

**Supplementary Figure 1: Expression of SMXL5-VENUS protein fusions in *smxl4;smxl5* mutants in comparison to promoter reporters.**

(**A – E**) Expression of SMXL5-VENUS protein fusions under the control of different heterologous promoters. Differentiated SEs are indicated by asterisks.

(**F**) Detection of the BRX-VENUS protein fusion expressed under the control of the *SMXL4* promoter. Differentiated SEs are indicated by the asterisk.

(**G – I**) Detection of *OPS:ER-VENUS*, *BAM3:ER-VENUS*, and *CVP2:ER-VENUS* reporter activities in *smxl4;smxl5* mutants. Arrow in G points to a weakly detectable reporter-derived signal.

(**J – M**) Detection of *APL:ER-VENUS* reporter activity in wild type (J, K) and *smxl4;smxl5* mutants (L, M). K and M show the same root tip as in J and L, respectively, without the counter stain signal. Scale bars in all pictures represent 50 µm. Arrows in J and K point to a weakly detectable reporter-derived signal.

(**N**) Root length measurements in different genetic backgrounds in the presence and absence of exogenous CLE45 five days after germination. Results from one representative experiment out of two independent experiments (n = 23–58 per experiment and genotype, except n = 6 for *bam3-2*) are shown. Mean values were analyzed by one-way ANOVA with post-hoc Tukey HSD (95 % CI). Statistical groups are indicated by letters.

**Supplementary Figure 2: Characterization of phloem development in *smxl* and *ops* mutants.**

(**A – K**) Phloem development is monitored in wild type (A), *smxl5* (B), *smxl4;smxl5* (C), *ops* (D-G) and *ops;smxl5* (H-K) mutant backgrounds by confocal analysis of mPS-PI-stained root tips.

Asterisks indicate the most apical appearance of differentiated SEs. In case ‘gaps’ are observed, more than one asterisk is depicted.

**Supplementary Figure 3: Characterization of phloem development in *smxl* and *brx* mutants.**

(**A – H**) Phloem development is monitored in wild type (A), *smxl5* (B), *smxl4;smxl5* (C), *brx* (D-E) and *brx;smxl5* (F-H) mutant backgrounds by confocal analysis of mPS-PI-stained root tips.

Asterisks indicate the most apical appearance of differentiated SEs. In case ‘gaps’ are observed, more than one asterisk is depicted.

**Supplementary Figure 4: *OBE3* cDNA clones isolated when screening for SMXL5 interactors using the Yeast-Two-Hybrid system.**

(A) Alignment of isolated clones (in green) to the full *OBE3* cDNA sequence (top) using CLC Main Workbench Version 7.6.1 (CLC Bio Qiagen, Aarhus, Denmark). Protein domains predicted for OBE3 are indicated above. The sequence area present in all isolated clones (selected interacting domain, SID) is marked by a purple arrow and flanked by red lines. The OBE3 protein contains two predicted nuclear localization signals (NLS) identified by the cNLS Mapper (Kosugi et al. 2009, NLS_bipartite and NLS_monopartite, light green arrows), a PHD- finger domain (blue arrow) with a histone 3 (H3) binding motif (light blue arrow) and a coiled coil domain (orange arrow) predicted by InterPro (EMBL-EBI, Cambridgeshire, UK). The OBE3 open reading frame is flanked by a 3’ and a 5’ untranslated region (3’UTR and 5’UTR, green arrows), respectively.

(B) Amino acid sequence encoded by the SID sequence, present in all isolated clones.

**Supplementary Figure 5: *SMXL4* or *SMXL5* genes do not genetically interact with *OBE1* or *OBE2* genes**

**(A)** 10 day-old *w*ild type, *smxl4, smxl5, smxl4;smxl5, obe1, obe2, smxl4;obe1, smxl4;obe2, smxl5;obe1, smxl5;obe2* seedlings are shown from left to right. Scale bar represents 1 cm.

**(B)** Quantification of root lengths depicted in A. Mean values of three independent experiments (n = 34-75 per experiment and genotype) were analyzed by a one-way ANOVA with post-hoc Tukey HSD (95 % CI). Statistical groups are indicated by letters.

**Supplementary Figure 6: SEs form in *smxl4* and *smxl5* single mutants**

(**A – D**) 2 day-old mPS-PI-stained root tips of wild type (A), *smxl4* (B), *smxl5* (C) and *smxl4;smxl5* (D) plants. Differentiating SEs were observed in wild type, *smxl4* and *smxl5* (pink arrows). Periclinal cell divisions are marked by orange and blue arrows. The QC is indicated by a yellow asterisk. Scale bars represent 20 µm.

(**E**) The distance of the first and second periclinal division from the QC for plants shown in A-D was quantified (n = 10-12). Statistical groups marked by letters were determined by one-way ANOVA with post-hoc Tamhane-T2 (95 % CI). Distances of 1^st^ cell divisions and 2^nd^ cell divisions were compared independently.

**Supplementary Figure 7: Activity of the *SMXL5:H4-GFP* transgene in wild type, *smxl5*, *smxl4;smxl5* and *smxl5;obe3* backgrounds and sorting of GFP-positive and GFP- negative nucleus fractions**

**(A)** Activity of the *SMXL5:H4-GFP* transgene in root tips of wild type, *smxl5*, *smxl4;smxl5* and *smxl5;obe3* seedlings two days after germination. Scale bars represent 20 µm.

**(B)** Sorting plots displaying gates for identifying GFP-positive (P4) and GFP-negative (P3) nuclei comparing wild type without transgene (wt ctrl) and wild type, *smxl5*, *smxl4;smxl5* and *smxl5;obe3* plants carrying the *SMXL5:H4-GFP* transgene. 10,000 sorting events were processed for display in all cases. P3 fractions included 2.2 to 2.9 % of all events and P4 fractions 0.1 to 0.2 % (0 % for wild type without the transgene). X-axes indicate RFP fluorescence (561 nm), Y-axes indicate GFP fluorescence (488 nm).

**Supplementary Figure 8: ATAC-seq analysis comparing GFP-positive and GFP-negative samples from wild type, *smxl5*, *smxl4;smxl5* and *smxl5;obe3* seedlings**

Profile of read alignment (dark and light green) and OCR detection (red) across all chromosomes for all samples processed. Dark green: GFP-positive samples. Light green: GFP-negative samples. Centromeric regions are indicated by reduced read alignment and OCR detection.

**Supplementary Figure 9: Chromatin conformation profile for *OPS* and *BAM3* gene regions.**

Profile of read alignment (dark and light green) and OCR detection (red) for all samples are shown. Dark green: GFP-positive samples. Light green: GFP-negative samples. Gene structures are indicated at the bottom (dark blue).

**Supplementary Figure 10: Chromatin conformation profile for *CVP2* and *APL* gene regions.**

Profile of read alignment (dark and light green) and OCR detection (red) for all samples are shown. Dark green: GFP-positive samples. Light green: GFP-negative samples. Gene structures are indicated at the bottom (dark blue).

**Supplementary Dataset 1: OCRs and associated genes specific for GFP-positive or GFP-negative fractions from wild type, *smxl5*, *smxl4;smxl5* or *smxl5;obe3* backgrounds, respectively.**

**Supplementary Datset 2: Genes with OCRs specific for GFP-positive or GFP-negative fractions for wild type, *smxl5*, *smxl4;smxl5* and *smxl5;obe3* backgrounds.**

**Supplementary Dataset 3: OCRs and associated genes specific for wild type, *smxl5*, *smxl4;smxl5* and *smxl5;obe3* plants comparing GFP-positive fractions from the respective backgrounds.**

**Supplementary Dataset 4: OCRs and associated genes specific for wild type, *smxl5*, *smxl4;smxl5* and *smxl5;obe3* plants comparing GFP-negative fractions from the respective backgrounds.**

**Supplementary Dataset 5: Phloem and non-phloem genes according to Brady et al., 2007.**

**Supplementary Table 1: Oligonucleotides used in this study.**

