## Supplementary Figure 1 for "OBERON3 and SUPPRESSOR OF MAX2 1-LIKE proteins form a regulatory module specifying phloem identity"

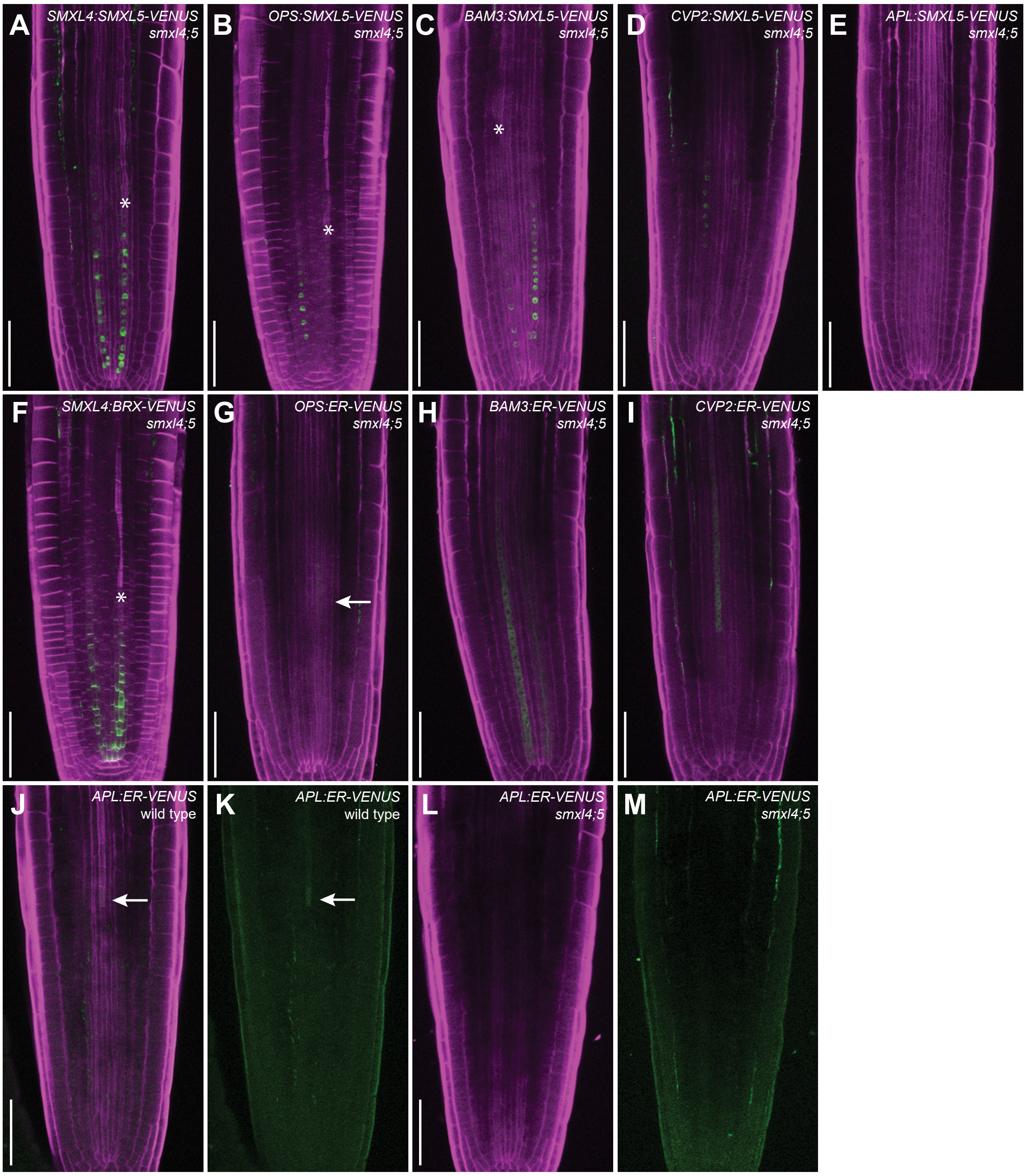


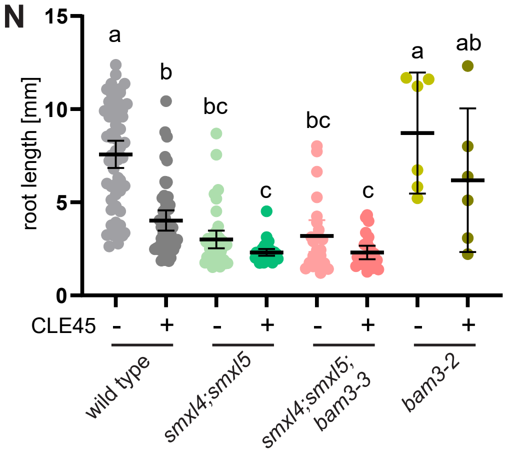


**Supplementary Figure 1: Expression of SMXL5-VENUS protein fusions in *smxl4;smxl5* mutants in comparison to promoter reporters.**

**(A – E)** Expression of SMXL5-VENUS protein fusions under the control of different heterologous promoters. Differentiated SEs are indicated by asterisks.

**(F)** Detection of the BRX-VENUS protein fusion expressed under the control of the *SMXL4* promoter. Differentiated SEs are indicated by the asterisk.

**(G – I)** Detection of *OPS:ER-VENUS*, *BAM3:ER-VENUS*, and *CVP2:ER-VENUS* reporter activities in *smxl4;smxl5* mutants. Arrow in G points to a weakly detectable reporter-derived signal.

**(J – M)** Detection of *APL:ER-VENUS* reporter activity in wild type (J, K) and *smxl4;smxl5* mutants (L, M). K and M show the same root tip as in J and L, respectively, without the counter stain signal. Scale bars in all pictures represent 50 µm. Arrows in J and K point to a weakly detectable reporter-derived signal.

**(N)** Root length measurements in different genetic backgrounds in the presence and absence of exogenous CLE45 five days after germination. Results from one representative experiment out of two independent experiments (n = 23–58 per experiment and genotype, except n = 6 for bam3-2) are shown. Mean values were analyzed by one-way ANOVA with post-hoc Tukey HSD (95 % CI). Statistical groups are indicated by letters.
