## Supplementary Figure 2 for "OBERON3 and SUPPRESSOR OF MAX2 1-LIKE proteins form a regulatory module specifying phloem identity"

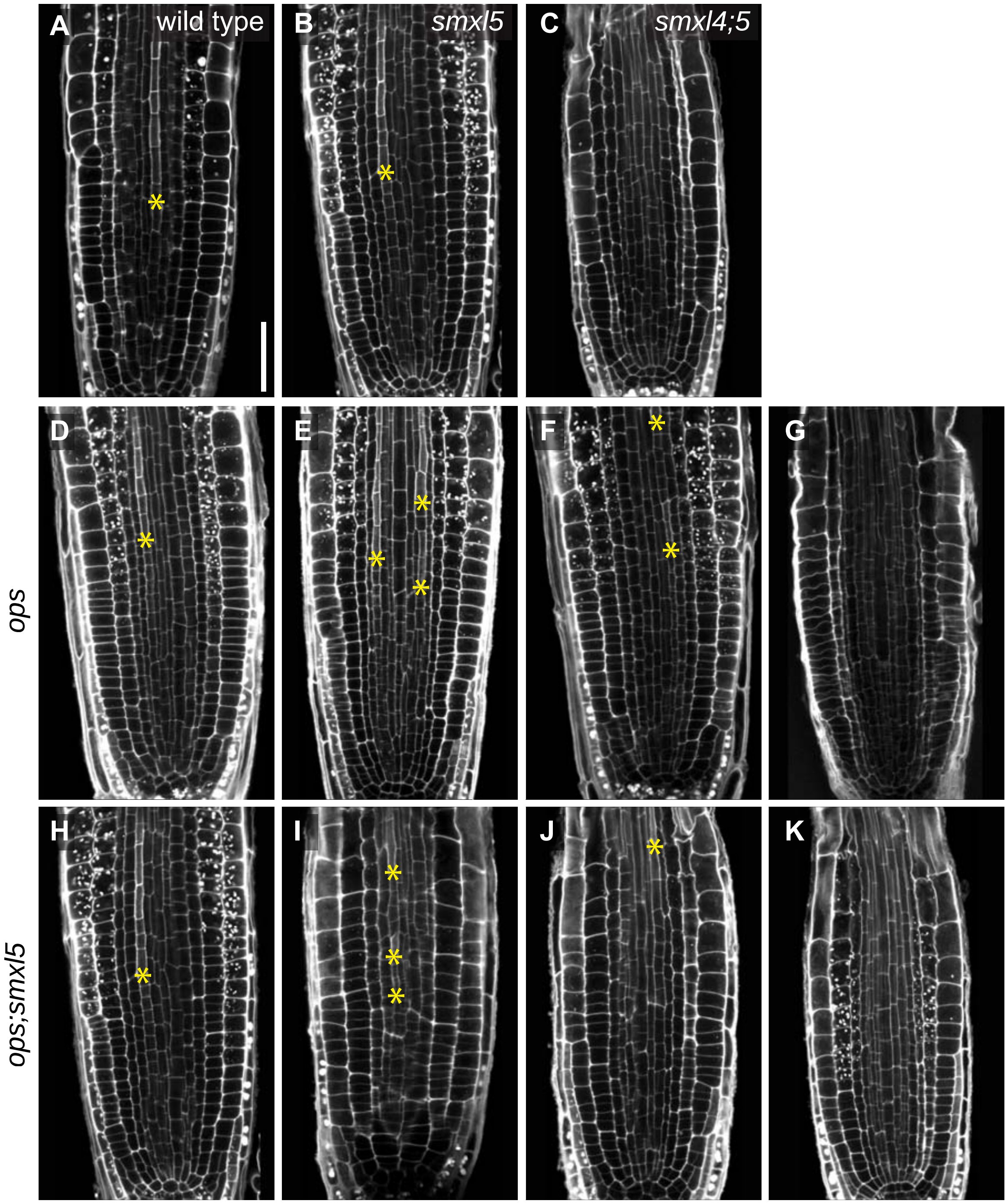


**Supplementary Figure 2: Characterization of phloem development in *smxl* and *ops* mutants.**
**(A – K)** Phloem development is monitored in wild type (A), *smxl5* (B), *smxl4;smxl5* (C), *ops* (D-G) and *ops;smxl5* (H-K) mutant backgrounds by confocal analysis of mPS-PI-stained root tips. Asterisks indicate the most apical appearance of differentiated SEs. In case 'gaps' are observed, more than one asterisk is depicted.
