## Supplementary Figure 4 for "OBERON3 and SUPPRESSOR OF MAX2 1-LIKE proteins form a regulatory module specifying phloem identity"

A

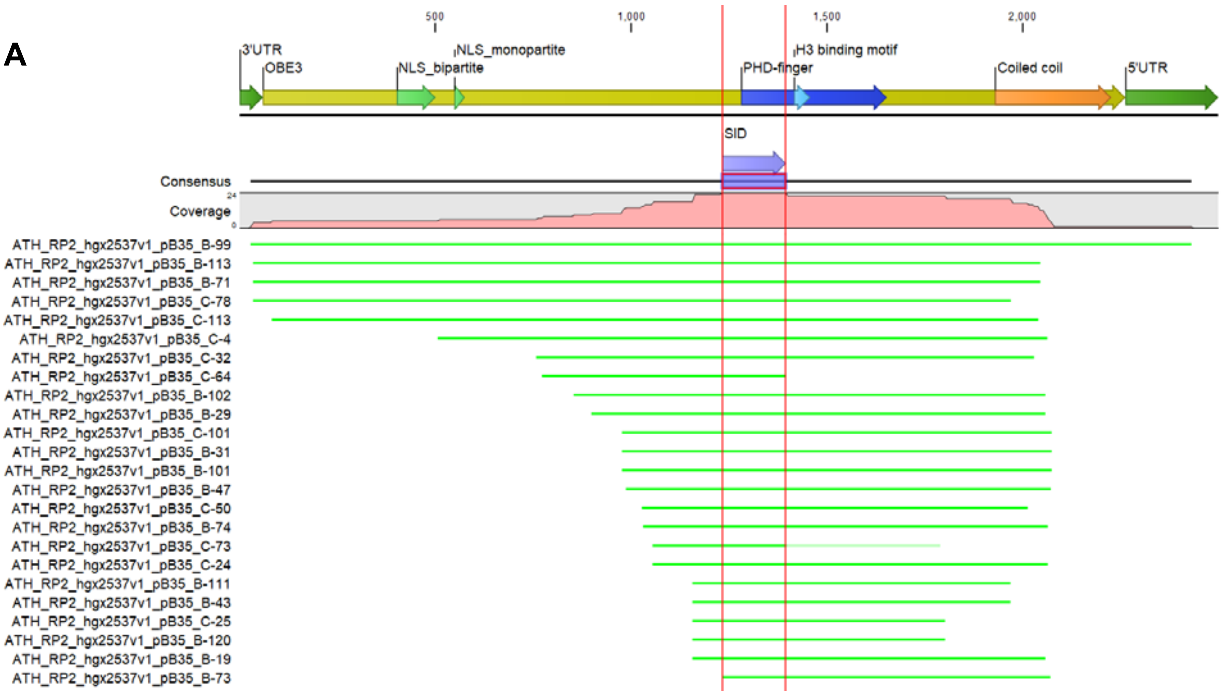


B

-IRIPMNELVEIFLFLRCRNVNCKSLLPVDDCECKICSNNKGFCSSCMCPVCLRF-

**Supplementary Figure 4: *OBE3* cDNA clones isolated when screeing for SMXL5 interactors using the Yeast-Two-Hybrid system.**

**(A)** Alignment of isolated clones (in green) to the full *OBE3* cDNA sequence (top) using CLC Main Workbench Version 7.6.1 (CLC Bio Qiagen, Aarhus, Denmark). Protein domains predicted for OBE3 are indicated above. The sequence area present in all isolated clones (selected interacting domain, SID) is marked by a purple arrow and flanked by red lines. The OBE3 protein contains two predicted nuclear localization signals (NLS) identified by the cNLS Mapper (Kosugi et al. 2009, NLS_bipartite and NLS_monopartite, light green arrows), a PHD-finger domain (blue arrow) with a histone 3 (H3) binding motif (light blue arrow) and a coiled coil domain (orange arrow) predicted by InterPro (EMBL-EBI, Cambridgeshire, UK). The OBE3 open reading frame is flanked by a 3' and a 5' untranslated region (3'UTR and 5'UTR, green arrows), respectively.

**(B)** Amino acid sequence encoded by the SID sequence, present in all isolated clones.
