## Supplementary Figure 5 for "OBERON3 and SUPPRESSOR OF MAX2 1-LIKE proteins form a regulatory module specifying phloem identity"

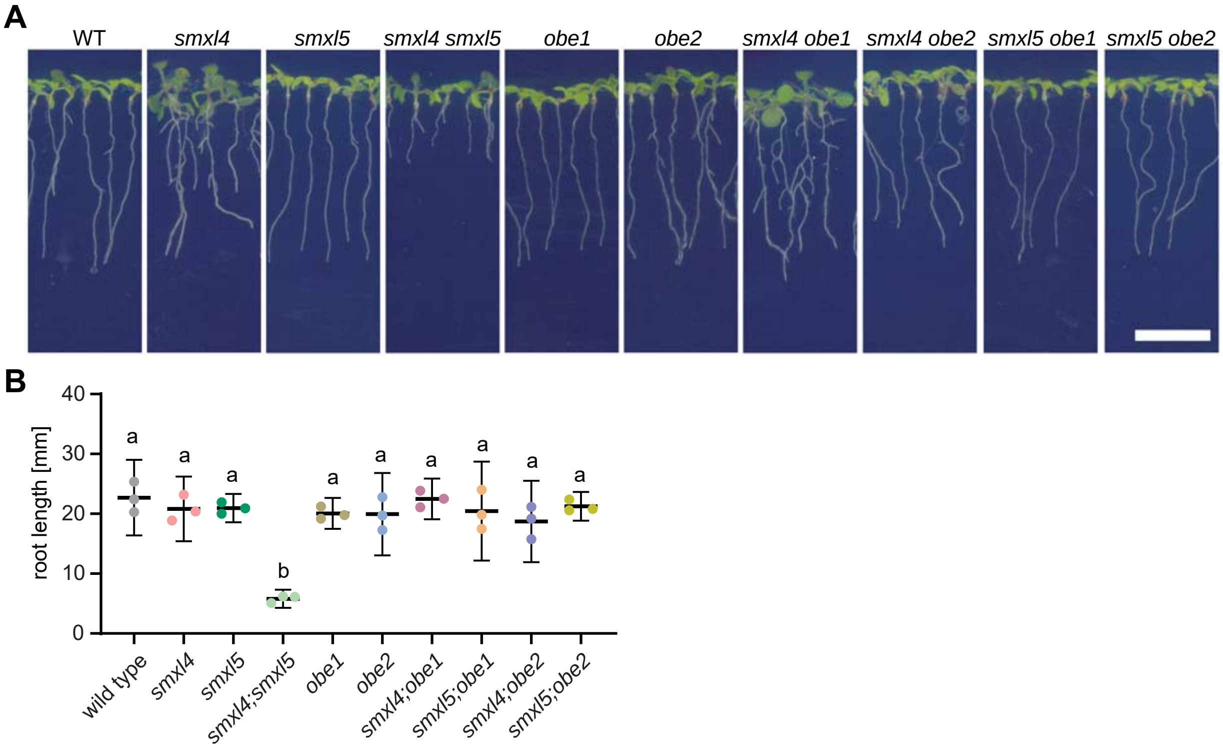


**Supplementary Figure 5: *SMXL4* or *SMXL5* genes do not genetically interact with *OBE1* or *OBE2* genes**
**(A)** 10 day-old wild type, *smxl4*, *smxl5*, *smxl4;smxl5*, *obe1*, *obe2*, *smxl4;obe1*, *smxl4;obe2*, *smxl5;obe1*, *smxl5;obe2* seedlings are shown from left to right. Scale bar represents 1 cm.
**(B)** Quantification of root lengths depicted in A. Mean values of three independent experiments (n = 34-75 per experiment and genotype) were analyzed by a one-way ANOVA with post-hoc Tukey HSD (95 % CI). Statistical groups are indicated by letters.
