## Supplementary Figure 6 for "OBERON3 and SUPPRESSOR OF MAX2 1-LIKE proteins form a regulatory module specifying phloem identity"

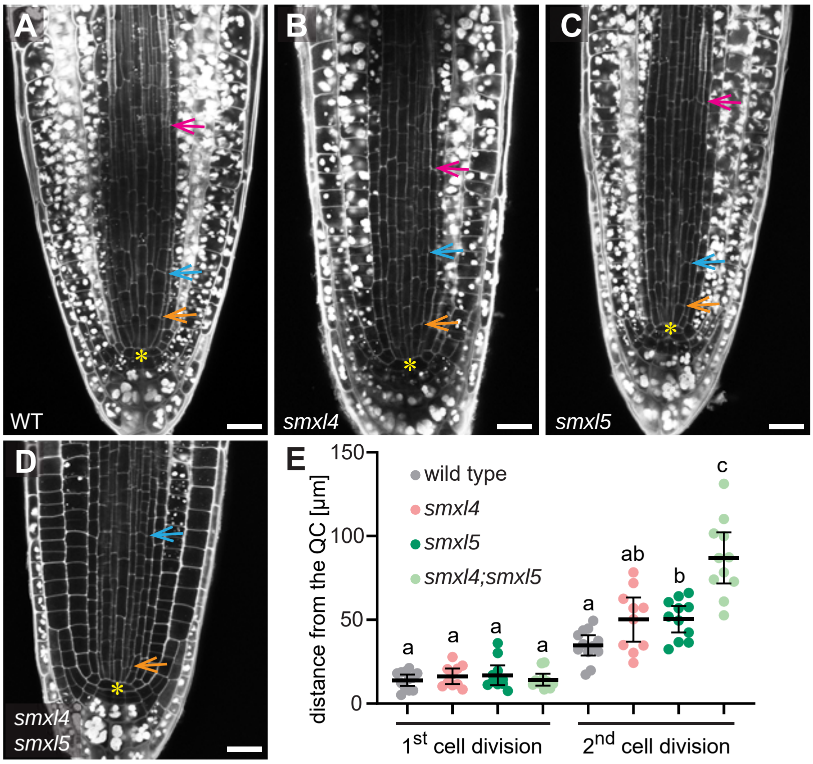


**Supplementary Figure 6: SEs form in *smxl4* and *smxl5* single mutants**

**(A – D)** 2 day-old mPS-PI-stained root tips of wild type (A), *smxl4* (B), *smxl5* (C) and *smxl4;smxl5* (D) plants. Differentiating SEs were observed in wild type, *smxl4* and *smxl5* (pink arrows). Tangential cell divisions are marked by orange and blue arrows. The QC is indicated by a yellow asterisk. Scale bars represent 20 µm.

**(E)** The distance of the first and second tangential division from the QC for plants shown in A-D was quantified (n = 10-12). Statistical groups marked by letters were determined by one-way ANOVA with post-hoc Tamhane-T2 (95 % CI). Distances of 1st cell divisions and 2nd cell divisions were compared independently.
