## Supplementary Figure 7 for "OBERON3 and SUPPRESSOR OF MAX2 1-LIKE proteins form a regulatory module specifying phloem identity"

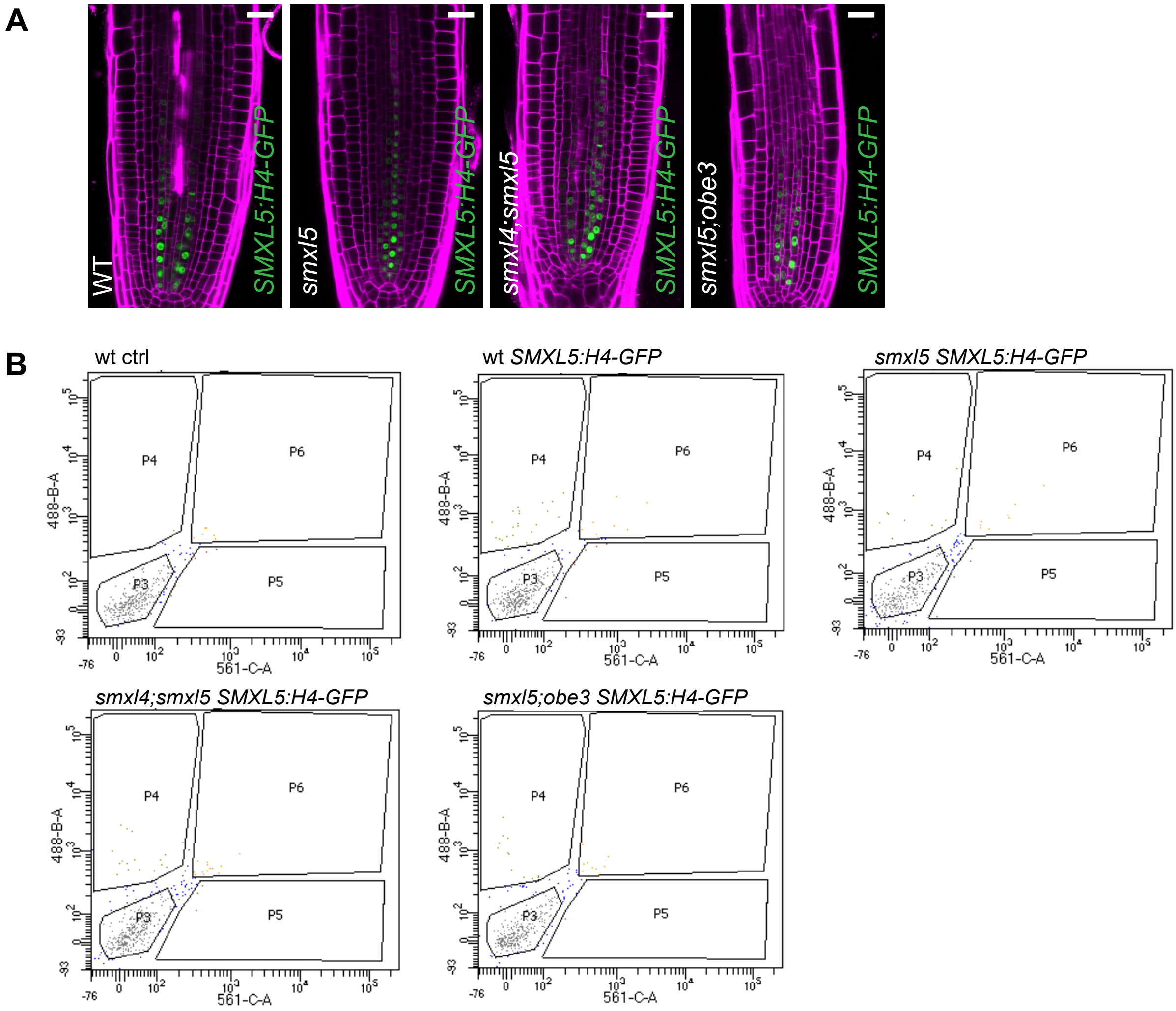


**Supplementary Figure 7: Activity of the *SMXL5:H4-GFP* transgene in wild type, *smxl5*, *smxl4;smxl5* and *smxl5;obe3* backgrounds and sorting of GFP-positive and GFP-negative nucleus fractions**

**(A)** Activity of the *SMXL5:H4-GFP* transgene in root tips of wild type, *smxl5*, *smxl4;smxl5* and *smxl5;obe3* seedlings two days after germination. Scale bars represent 20 µm.

**(B)** Sorting plots displaying gates for identifying GFP-positive (P4) and GFP-negative (P3) nuclei comparing wild type without transgene (wt ctrl) and wild type, *smxl5*, *smxl4;smxl5* and *smxl5;obe3* plants carrying the *SMXL5:H4-GFP* transgene. 10,000 sorting events were processed for display in all cases. P3 fractions included 2.2 to 2.9 % of all events and P4 fractions 0.1 to 0.2 % (0 % for wild type without the transgene). X-axes indicate RFP fluorescence (561 nm), Y-axes indicate GFP fluorescence (488 nm).
