## Supplementary Figure 8 for "OBERON3 and SUPPRESSOR OF MAX2 1-LIKE proteins form a regulatory module specifying phloem identity"

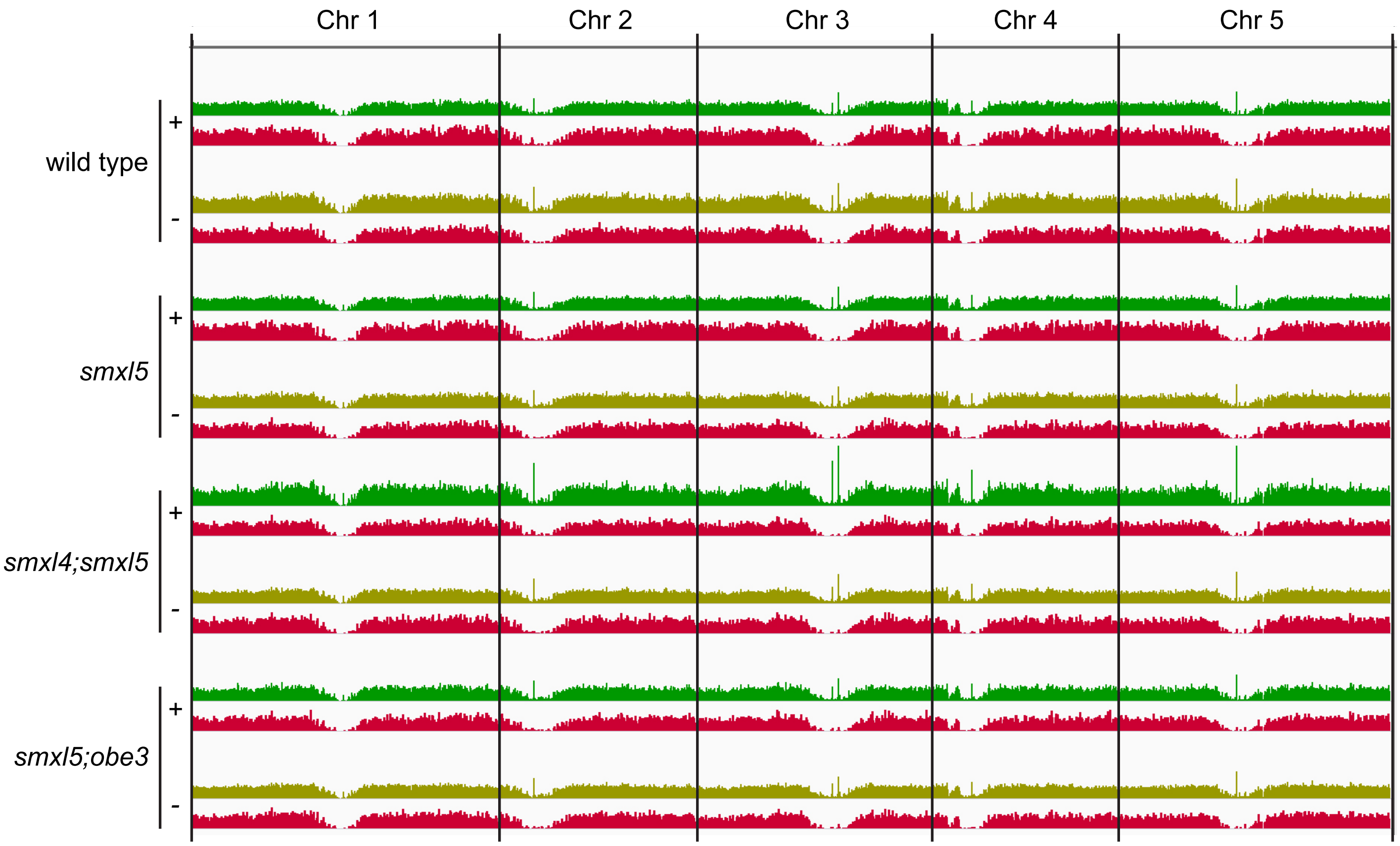


**Supplementary Figure 8: ATAC-seq analysis comparing GFP-positive and GFP-negative samples from wild type, *smxl5*, *smxl4;smxl5* and *smxl5;obe3* seedlings**

Profile of read alignment (dark and light green) and OCR detection (red) across all chromosomes for all samples processed. Dark green: GFP-positive samples. Light green: GFP-negative samples. Centromeric regions are indicated by reduced read alignment and OCR detection.
