## Supplementary Figure 9 for "OBERON3 and SUPPRESSOR OF MAX2 1-LIKE proteins form a regulatory module specifying phloem identity"

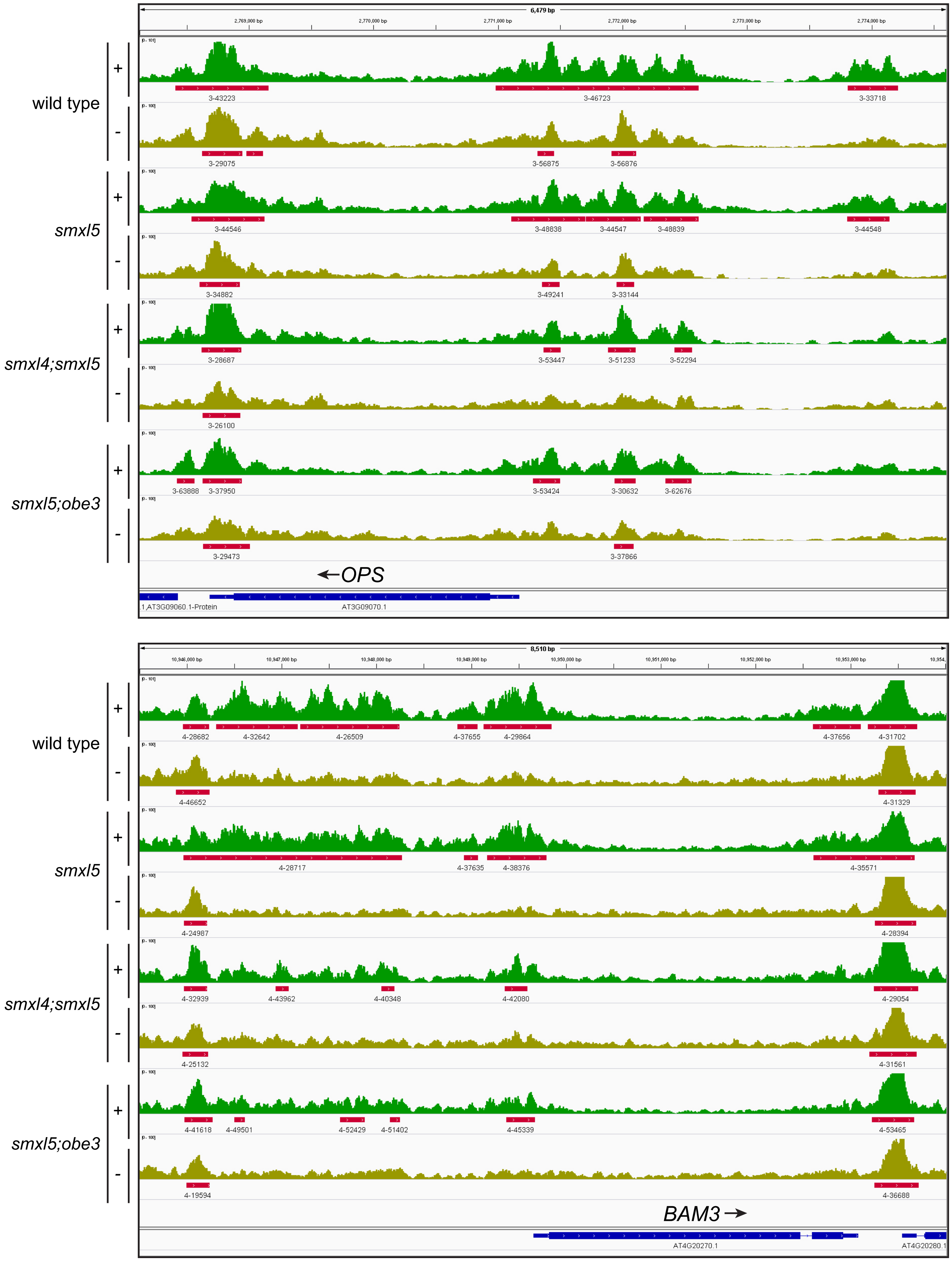


**Supplementary Figure 9: Chromatin conformation profile for *OPS* and *BAM3* gene regions.**
Profile of read alignment (dark and light green) and OCR detection (red) for all samples are shown. Dark green: GFP-positive samples. Light green: GFP-negative samples. Gene structures are indicated at the bottom (dark blue).
